# Modeling motor-evoked potentials from neural field simulations of transcranial magnetic stimulation

**DOI:** 10.1101/847830

**Authors:** Marcus T Wilson, Bahar Moezzi, Nigel C Rogasch

## Abstract

**Objective:** To develop a population-based biophysical model of motor-evoked potentials (MEPs) following transcranial magnetic stimulation (TMS).

**Methods:** We combined an existing MEP model with population-based cortical modeling. Layer 2/3 excitatory and inhibitory neural populations, modeled with neural-field theory, are stimulated with TMS and feed layer 5 corticospinal neurons, which also couple directly but weakly to the TMS pulse. The layer 5 output controls mean motoneuron responses, which generate a series of single motor-unit action potentials that are summed to estimate a MEP.

**Results:** A MEP waveform was generated comparable to those observed experimentally. The model captured TMS phenomena including a sigmoidal input-output curve, common paired pulse effects (short interval intracortical inhibition, intracortical facilitation, long interval intracortical inhibition) including responses to pharmacological interventions, and a cortical silent period. Changes in MEP amplitude following theta burst paradigms were observed including variability in outcome direction.

**Conclusions:** The model reproduces effects seen in common TMS paradigms.

**Significance:** The model allows population-based modeling of changes in cortical dynamics due to TMS protocols to be assessed in terms of changes in MEPs, thus allowing a clear comparison between population-based modeling predictions and typical experimental outcome measures.

**Highlights:**

- A model of motor-evoked potential formation gives a realistic electromyogram in response to TMS.
- The model reproduces effects of SICI, ICF and LICI.
- A link between existing neural field modeling and realistic outcome measures of TMS is provided.

## 1. Introduction

Transcranial Magnetic Stimulation (TMS) is a non-invasive form of brain stimulation used for the study of brain function and for clinical treatments of brain disorders such as depression (Hallett, 2007; Ziemann et al., 2008; Pascual-Leone et al., 2000; Lefaucheur et al., 2014). Applying a single TMS pulse at sufficient intensity over the primary motor cortex results in firing of layer 5 corticospinal neurons due mainly to transsynaptic activation from layer 2/3 interneurons and horizontal fibres, but also direct activation of the neurons by the pulse at high enough intensities (Hallett, 2007; Di Lazzaro et al., 2012). The descending volley of activity gives a measurable motor response in peripheral muscles targeted by the stimulated region, known as a motor-evoked potential (MEP) (Hallett, 2000). MEPs have been widely used as a measure of the excitability of the corticomotor system in TMS studies, and have revealed several well known neural phenomena related to TMS, such as periods of net inhibition and excitation using paired pulse protocols (i.e. short and long interval intracortical inhibition [SICI; LICI], and intracortical facilitation [ICF]) (Valls-Solé et al., 1992; Kujirai et al., 1993), and a cortical silent period observed when TMS is given during a voluntary contraction. Furthermore, MEPs are used to assess changes in cortical excitability resulting from repetitive TMS (rTMS) protocols (Di Lazzaro et al., 2008), which are thought to induce plasticity in cortical circuits through mechanisms similar to long-term potentiation and depression (LTP/D) (Cooke and Bliss, 2006). However, despite nearly 30 years of research, it remains unclear how microscale mechanisms underlying plasticity occurring at synaptic level (e.g. LTP/D) manifest when large populations of neurons are activated as with TMS (Parkin et al., 2015; Matheson et al., 2016).

Biophysically-informed models provide a mathematical description of TMS and other neurostimulation effects that can be used to better understand TMS phenomena (Seo and Jun, 2017; Wilson et al., 2018). Models typically describe biophysical processes with equations. Existing models include descriptions of the shape and timecourse of the magnetic and induced electric fields due to TMS, including realistic human head geometries (Thielscher et al., 2011; Deng et al., 2013; Opitz et al., 2013; Tang et al., 2016; Bungert et al., 2016), large networks of spiking neurons (Esser et al., 2005), detailed descriptions of spiking of single neurons and small networks of neurons (Traub et al., 2003; Rusu et al., 2014; Moezzi et al., 2017), population-based descriptions of neural firing rates (Deco et al., 2008; Pinotsis et al., 2014) and plasticity effects (Fung et al., 2013; Wilson et al., 2016).

Modeling of the processes underlying TMS-induced effects has been undertaken through several stategies (Seo and Jun, 2017; Wilson et al., 2018). Esser et al. (2005) have constructed a low-level model of 33 thousand neurons in the cortex and thalamus with five million synaptic connections. The model demonstrates biologically-plausible spontaneous activity and evoked responses, notably I-waves. More detailed multicompartment models for single neurons or small groups of neurons have also been used, to study bursting phenomena in more detail (Traub et al., 2003). Rusu et al. (2014) used a detailed model of a layer 5 neuron, fed by a small population of layer 2/3 single-compartment cells. The hypothesis is that interactions between layer 2/3 and layer 5 cells are purely feed-forward (no resonance loops or chains of excitatory and inhibitory cells), and a spike generated in the layer 2/3 cells affects the layer 5 cell after a certain time (Triesch et al., 2015). Under this hypothesis, I-waves could be reproduced. However, the mechanism for generation of the I-waves in this model differs from the more conventional view that I-waves are a result of repetitive input to layer 5 cells from a resonanting circuit.

While models of individual neurons have proven useful in capturing TMS-related phenomena, there are several notable limitations. First, the high number of parameters (e.g. values determining receptor conductances, synapse weights, conduction delays etc. for each neuron/synapse) can pose problems in constraining the model, thereby increasing the risk of over-fitting. Second, the complexity of these models often comes with a high computational cost, which greatly increases the time required for simulations. Finally, the models have largely focused on generating corticospinal output which is suitable for capturing I-wave activity, but does not generalise to MEPs measured at peripheral muscles, which is by far the most common experimental method for assessing TMS-evoked activity of the motor system.

While the dynamics of evoked responses in the brain involves highly non-linear processes, modeling these need not be complicated. Population-based modeling (Deco et al., 2008; Pinotsis et al., 2014), including neural mass or neural field approaches, considers firing rates of populations of cells, rather than detailed dynamics of many individual cells. As such, population-based models have far fewer parameters, and are less computationally expensive than models of individual neurons. Neural field modeling is well suited to TMS because a TMS pulse excites many thousands of neurons over an area of several centimeters-squared. Neural field approaches have been used to model cortical plasticity following repetitive TMS, a lasting change in strengths of connections between neurons (Huang et al., 2011; Fung et al., 2013; Fung and Robinson, 2014; Wilson et al., 2016). In these works, plasticity has been included using rules which capture either phenomenological descriptions of plasticity (e.g. spike timing dependent plasticity), or physiological theories (e.g. calcium dependent plasticity) (Shouval et al., 2002).

Population-based cortical modeling of TMS has been hard to interpret in relation to human experiments. For example, most models have evaluated changes in synaptic weights between excitatory neural populations following rTMS but it remains unclear how these changes would impact the amplitude of MEPs. To address this failing in the existing literature, population based models must be combined with models of the motor system.

Li et al. (2012) have described MEP amplitude and shape in terms of a sum of individual motor unit responses, with thresholds for the motor units distributed exponentially. Such an approach has the benefit of simplicity, with few parameters. However, it describes only the motoneuron response, not the processes that feed it, and misses biophysical detail such as motor unit synchrony. Moezzi et al. (2017) have developed this further; they have used the hypothesis and I-wave model of Rusu et al. (2014) to simulate MEP formation following TMS using a population of layer 2/3 excitatory and inhibitory neurons, feeding layer 5 cortical cells and motoneurons. They reproduced MEP responses that matched closely those measured experimentally; however, this was under similar limitations to other individual neuron models in terms of a large parameter space and high computational cost. In contrast, Goetz et al. (2019) have used a statistical model based on experimental data to compile a MEP model, although this model was not biophysical in the sense that it only modeled the distribution of MEPs, not the biological processes underlying MEP generation.

The aim of this study was to develop an MEP model that can be used alongside a general population-based neural field model of single-pulse, paired-pulse and repetitive TMS, thereby linking population-based model predictions of the effects of TMS with commonly-used experimental outcomes for the first time. To achieve this aim, we have combined the approach of Moezzi et al. (2017) with neural field modeling. Specifically, we model the layer 2/3 and layer 5 populations with population-based dynamics. Firing rates of motoneurons are described as functions of the layer 5 firing rate, and a train of motoneuron firings is reconstructed. Each motounit contributes a motounit action potential (MUAP). Thus MEP activity is determined. This approach provides a much-needed link between population-based models of cortical dynamics, and models of MEP activity, thereby allowing a direct comparison between model outputs and human experiments. To test the generalizability of our MEP model, we first assess whether we can capture well known single and paired-pulse MEP phenomena. We then evaluate how sensitive our MEP model is to changes in synaptic weights predicted by population-based rTMS models of plasticity.

## 2. Methods

We have combined a neural field approach (Fung and Robinson, 2014; Wilson et al., 2016) with existing models of MEP formation (Li et al., 2012; Moezzi et al., 2017). The scheme is shown schematically in Fig. 1.

**Figure 1:**
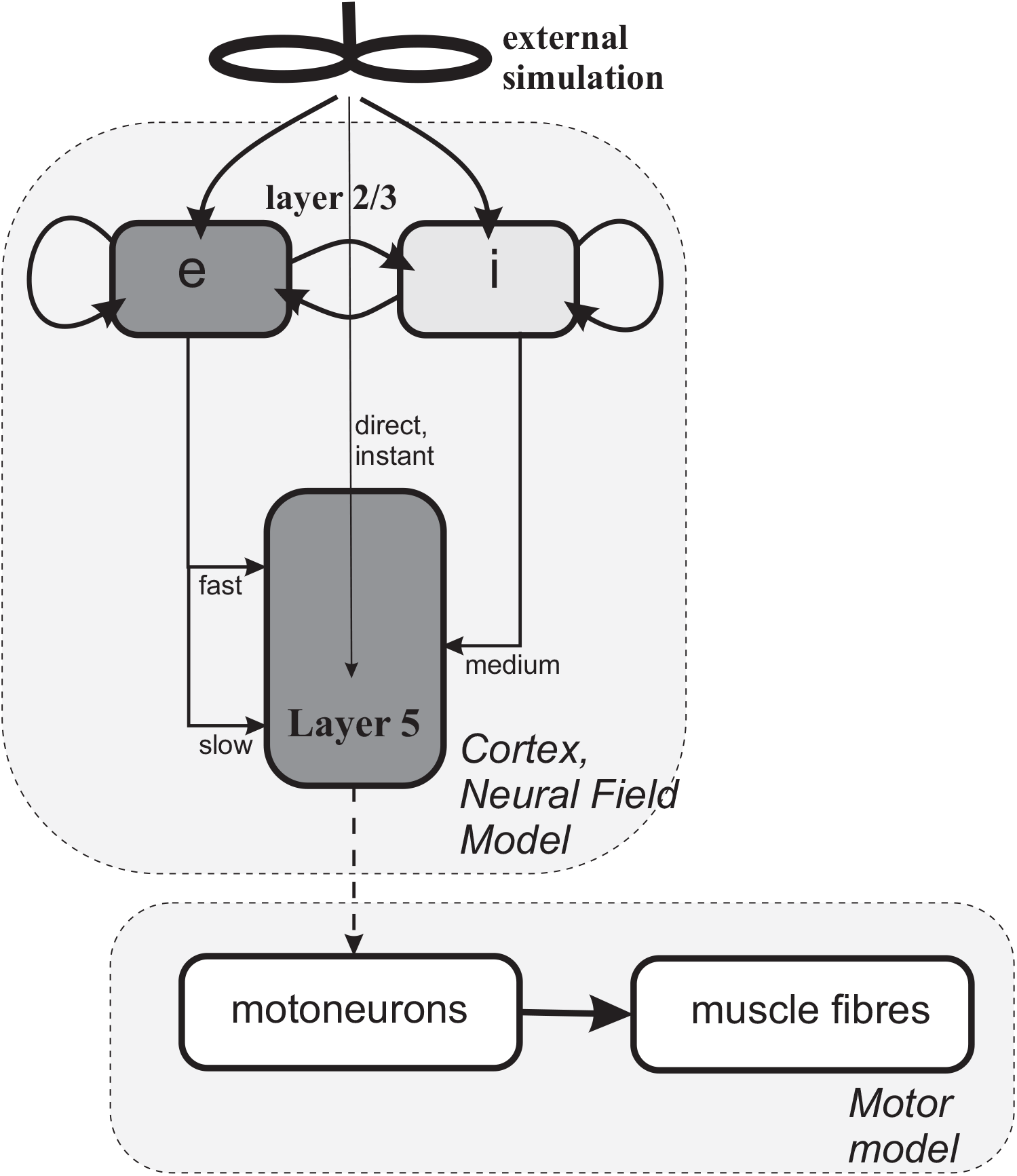
A schematic of the modeling approach. Transcranial magnetic stimulation (TMS) stimulates populations of layer 2/3 excitatory and inhibitory neurons. The layer 2/3 populations stimulate each other. They have multiple projections to a population of layer 5 excitatory corticospinal output neurons which also receive low-intensity direct stimulation from the TMS pulses. These three are modeled with NFTsim (Sanz-Leon et al., 2017). The axonal flux from the layer 5 corticospinal output neurons stimulates a population of motoneurons within the spinal cord; each motoneuron firing produces a response from the muscle fibres. The sum of these fibre responses produces a motor-evoked potential (MEP). The latter components are modeled with the approach of Li et al. (2017).

### 2.1. Neural Field components (Cortical model)

Neural Field Theory (NFT) provides a nonlinear, statistical model for the dynamics of populations of neuronal cells and their interactions via dendrites and axons (Deco et al., 2008; Pinotsis et al., 2014; Wright and Liley, 1996; Robinson et al., 1997; Breakspear, 2017). Population-averaged properties such as mean firing rate and axonal pulse rate are modeled as a function of time *t*. We have used the NFTsim model (Sanz-Leon et al., 2017). The mathematical description and parameter values are summarized in Appendix A.

Specifically, a population of layer 2/3 excitatory neurons (labeled ‘*e*’), a population of layer 2/3 inhibitory neurons (*i*), and a population of layer 5 excitatory corticospinal output neurons (*v*) are modeled with NFTsim. These populations couple together as indicated by the arrows in Fig. 1. They are also coupled to an external driving population (labeled *x*) describing the TMS application. Coupling strengths *to* population *a from* population *b* are denoted in this paper by *ν_ab_*, where *a* and *b* can take the labels *e*, *i*, *x* or *v*. Synaptic responses include excitatory receptors and both fast-acting *γ*-aminobutyric acid A (GABA_A_) and slower-acting *γ*-aminobutyric acid B (GABA_B_) receptor effects (Wilson et al., 2016, 2018).

The layer 2/3 populations project to a population of layer 5 excitatory corticospinal output neurons (*v*). There are multiple projections to different parts of the dendritic tree (Moezzi et al., 2017). Excitatory connections are made with short and long propagation delays (specifically 1 ms and 5 ms respectively), to model connections to basal and apical dendrites of the cortiospinal neurons (Petreanu et al., 2009). Inhibitory connections are made with a medium time delay (specifically 3 ms), to align with the hypothesis of Rusu et al. (2014).

The layer 2/3 excitatory, layer 2/3 inhibitory and layer 5 populations plus the external stimulation are modeled with NFTsim (Sanz-Leon et al., 2017). Although there are many parameters, many have physical constraints placed on them (Robinson et al., 2004). Parameters for the layer 2 and 3 cells have been chosen to be consistent with previous modeling (Wilson et al., 2016, 2014); for the layer 5 cells the firing response to synaptic input has been tuned to give plausible responses to stimulation, broadly consistent with Moezzi et al. (2017), including maximum population firing rate of 300 s^−1^ and a rapid climb in output once threshold has been reached at a mid-range stimulation intensity.

The NFT modeling gives the mean axonal flux rate of the layer 5 neural population as a function of time. This firing rate is then used as an input (dotted arrow in Fig. 1) to the next stage of modeling, summarized by the lower gray box in the figure.

TMS stimulation of the cortex can lead to both direct (D-) and indirect (I-) waves of descending activity, recorded in the epidural space (Hallett, 2007; Di Lazzaro et al., 2012). Often several indirect waves are recorded, at several hundred hertz frequency. While their origin has not been precisely established, they are likely to be a result of TMS-induced activity within the cortex propagating down nerve pathways. Rusu et al. (2014) presented a simple model of I-wave formation by projecting populations of layer 2 and 3 cells onto a compartmentalized layer 5 neuron. Averaged responses of many cases showed synchrony in layer 5 firings, resulting in I-waves of activity. Moezzi et al. (2017) have demonstrated similar synchrony by modeling explicitly many layer 5 cells simultaneously. In our model it is not possible to capture I-waves in a similar way, since neural synchrony is not explicitly captured when only population-averaged rates are considered because exact timings of firings are not explicitly modeled (Wilson et al., 2018, 2012). That is, a mean firing rate of a population does not tell us about the synchrony of firings within the population. While the population approach is not well-suited to capturing firing events such as those generating I-waves, it is well suited for capturing the slower shifts in net excitation and inhibition which are thought to underlie paired-pulse phenomena such as SICI and ICF.

### 2.2. Modeling TMS inputs

A particularly challenging aspect of developing this model was deciding how to model TMS inputs to the cortical populations. There is little direct evidence on how TMS interacts with specific neuronal populations within the cortex in humans. Indirect evidence from surface electromyography and spinal epidural recordings suggest that layer 5 corticospinal output neurons (CSNs) are not directly activated by TMS at subthreshold and low/moderate suprathreshold intensities. Instead, CSNs are likely activated transynaptically by interneuron populations presumably located in layers 2/3, as well as long-range horizontal connections from other cortical regions which are preferentially activated by the TMS pulse (Di Lazzaro and Rothwell, 2014). Transynaptic activation of CSNs is further supported by invasive recordings in rodents, showing preferential activation of neurons in superficial layers by TMS without activation of layer 5 neurons at moderate intensities (Murphy et al., 2016). At subthreshold intensities, paired pulse studies in humans show a lower threshold for MEP inhibition than facilitation (Ziemann et al., 1996), suggesting that inhibitory interneurons have the lowest activation threshold by TMS, although this may result from stimulation of upstream excitatory interneurons which synapse onto the inhibitory interneurons (Di Lazzaro and Rothwell, 2014). Direct activation of CSNs can occur at higher suprathreshold intensities (e.g. >150% RMT), particularly for latero-medial coil orientations (Di Lazzaro et al., 1998), although this appears more difficult using posterior-anterior orientations which are more typically used in TMS studies. Aside from stimulation intensity and coil orientation, TMS activation patterns are also altered by different coil types, pulse widths and pulse shapes (Di Lazzaro and Rothwell, 2014), demonstrating the complexity of TMS-cortical interactions.

Most modeling approaches to date have focused on estimating the spatial distribution of the electric field generated by TMS in the gray and white matter (Laakso et al., 2018; Bungert et al., 2016; Opitz et al., 2011), however several studies have attempted to further refine this approach by modeling how these fields interact with morphologically-realistic neurons within the cortex (Pashut et al., 2014; Seo and Jun, 2017; Aberra et al., 2020). The most detailed of these models found the lowest activation thresholds in layer 2-5 neurons with pyramidal neurons and inhibitory interneurons showing similar activation thresholds (Aberra et al., 2020), although this model did not include CSNs. Furthermore, it remains unclear from these static anatomical models how activation by TMS would alter ongoing firing rates within these different neuronal populations across different stimulation intensities.

Given the complex nature of estimating how TMS interacts with cortical circuits, we have adopted a pragmatic approach. In NFTsim, stimulation is applied with an ‘external’ rate *ϕ*_*x*_(*t*) (Fung et al., 2013), which can be interpreted as average number of action potentials per second that are introduced along each axon in the cortical populations. For example, a stimulus intensity of 1000 s^−1^ for 0.5 ms introduces on average 0.5 action potential onto each axon (e.g. 50% of neurons activated). Invasive studies recording individual cortical neurons following TMS in primates found that 28% of neurons changed firing rate following stimulation at 120% of RMT, giving a stimulus intensity of order 600 s^−1^ for 0.5 ms as a broad estimate for generating moderate size MEPs. We assume increasing external rate *ϕ*_*x*_ corresponds to an increasing TMS machine output, though the linearity of this relationship has not been established.

Based on experimental findings in humans (Ziemann et al., 1996; Ilic et al., 2002), we make the assumption that the strength of this drive to different neural populations depends upon the TMS intensity, with low intensities preferentially stimulating the inhibitory neurons over the excitatory neurons in layer 2/3, but higher intensities strongly stimulating the excitatory neurons, possibly due to the geometry of the axonal connections (Silva et al., 2008). While plausible, we acknowledge there is little direct evidence for this. Furthermore, we assume layer 5 CSNs receive lower direct inputs from TMS than layer 2/3 neurons. Hence, the external TMS to layer 2/3 excitatory coupling *ν*_*ex*_ is modeled as a function of stimulus intensity *ϕ*_*x*_ with a sigmoid relationship:

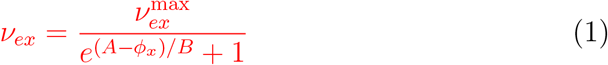

where 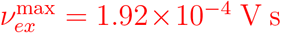 is the maximum external to layer 2/3 excitatory coupling, with *A* = 500 s^−1^ and *B* = 100 s^−1^ describing the threshold and width of the curve respectively. The external TMS to inhibitory cell coupling *ν*_*ix*_ is modeled as a constant:

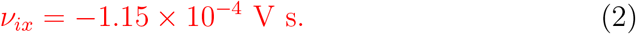

We use a low strength for the layer 5 external coupling compared to the layer 2/3 external coupling; specifically we set *ν*_*vx*_ = 0.1*ν*_*ex*_:

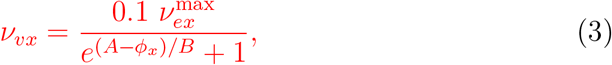

where 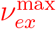, *A* and *B* are as defined above. Changes in external coupling to TMS across intensities within the different populations are plotted in Fig. 2. We emphasize that such preferential stimulation is an assumption in our modeling with only indirect experimental evidence, but we show the effects of varying the *ν*_*ix*_ and Eq. (1) parameters in Section 3.

**Figure 2:**
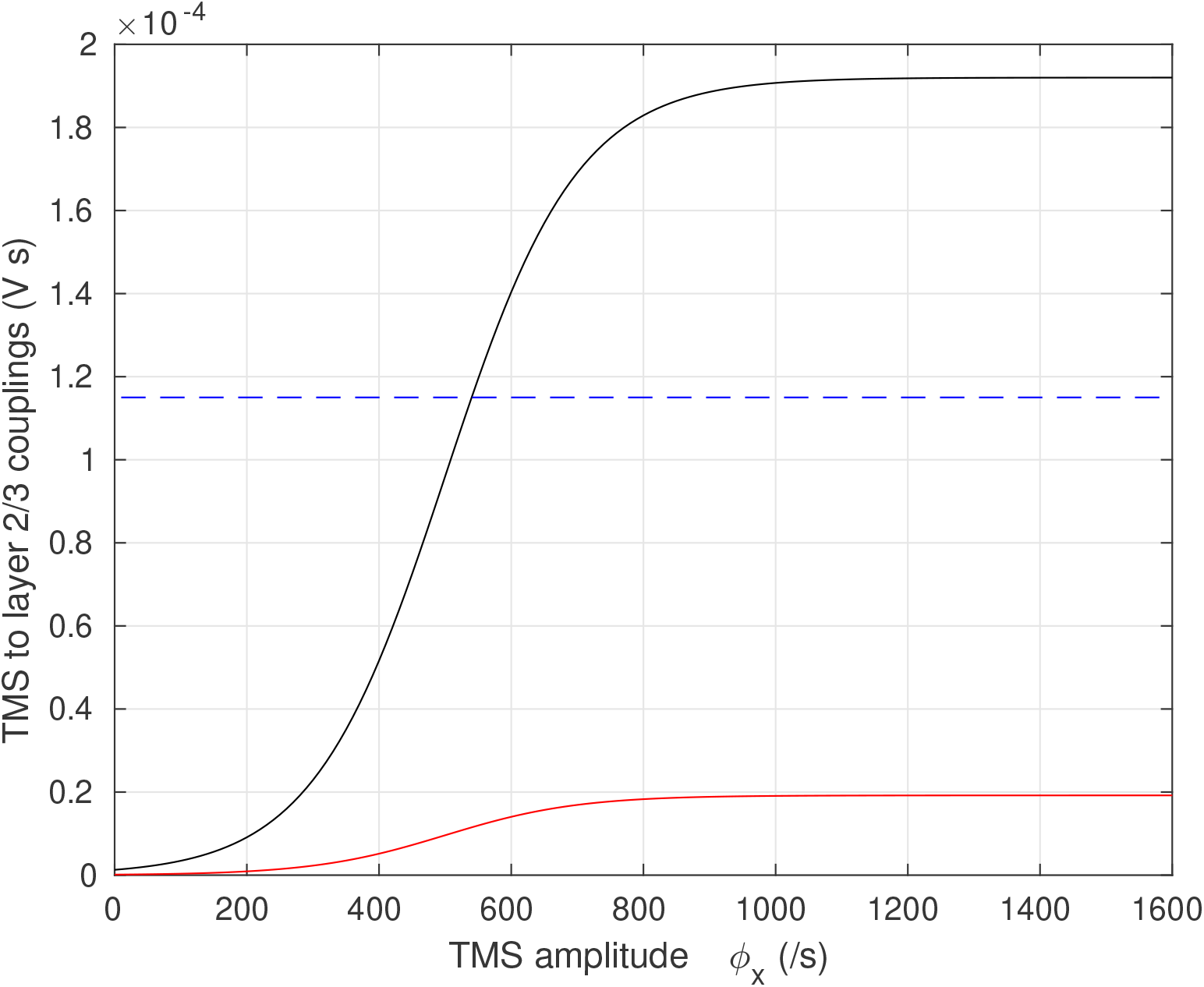
The external TMS to excitatory and inhibitory couplings as a function of external rate *ϕ*_*x*_ (assumed to be proportional to TMS amplitude). The TMS-to-excitatory coupling *ν*_*ex*_, given by Eq. (1), is shown by the solid black line; the TMS-to-inhibitory coupling *ν*_*ix*_ is shown by the dashed blue line. The TMS-to-layer 5 corticospinal output neuron coupling is shown by the solid red line.

### 2.3. MEP model components

Population-averaged cortical responses are often linear or nearly-linear, making NFT appropriate (Deco et al., 2008; Wilson et al., 2012). However, motor responses are challenging to describe in a linear way; for example 20 motoneurons firing at 100 s^−1^ produces a very different MEP to 10 motoneurons firing at 200 s^−1^.

Instead, we use a model of Li et al. (2012) to describe MEP formation. Here, the layer 5 axonal pulse rate *ϕ*_*v*_ is used to determine the rate of firing of *N* = 100 motoneurons. Each motoneuron, indexed by *k*, has an instantaneous firing rate *Q*_*k*_ (*k* = 1 … *N*) that is a function of the axonal flux rate from the layer 5 cells (Li et al., 2012), that is *Q*_*k*_ = *f*_*k*_(*ϕ*_*v*_).

All motoneurons have a threshold input below which they do not fire. Thresholds are distributed exponentially:

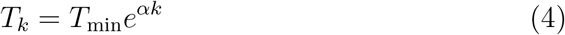

where *T*_*k*_ is the threshold of the *k*-th unit and *T*_*min*_ is a minimum threshold (set to 14 s^−1^ (Li et al., 2012)). The parameter *α* is set through

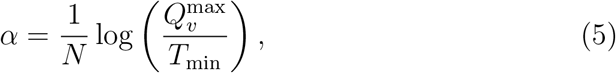

where 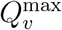 is the maximum layer 5 firing rate, so that the final (*N*-th) motor unit has a threshold corresponding to the maximum layer 5 firing rate. When the axonal flux rate *ϕ*_*v*_ from the layer 5 neurons exceeds a unit’s threshold *T*_*k*_, the unit fires with a rate given by the following function of axonal flux:

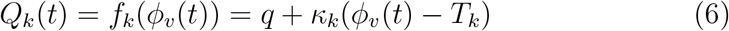

where *q* is the minimum firing rate (which we set at 8 Hz for all units) and *κ*_*k*_ is a constant for each *k*. The gradient *κ*_*k*_ is chosen separately for each *k* so that all motoneurons reach the same *Q*_*k*_, equal to 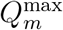 when *ϕ*_*v*_ is equal to 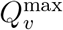, the maximum firing rate of the layer 5 neurons. We set 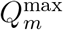 to 300 s^−1^ and 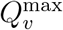, to 900 s^−1^, to broadly align with previous simulations at high (150% resting motor threshold, RMT) pulse intensity (Moezzi et al., 2017).

We next determine the times at which the motoneurons fire. We denote by 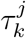, where *j* and *k* are both positive integers, the *j*-th firing event of the *k*-th unit. To identify these events we time-integrate the firing rates *Q*_*k*_(*t*) (Wilson et al., 2012). When the time-integral of the firing rates between two times is equal to 1, we can say that on average each neuron in the population has a single firing event between these two times. Thus, we identify the *j*-th firing event when the time-integral of the firing rate reaches *j*. That is, the 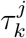 are the times such that:

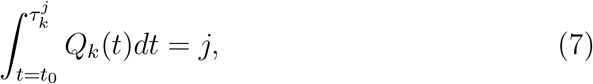

where *t*_0_ is the time that the firing rate of the neuron crosses its threshold value.

We produce a MEP by summing individual motounit action potentials (MUAPs). Each motoneuron leads to a MUAP whose size *M*_*k*_ is proportional to its threshold — that is, the units recruited latest fire the strongest. Thus *M*_*k*_ = *M*_0_*e*^*αk*^, where *M*_0_ is a constant set to be 42 mV s^−1^ so that the MEP’s amplitude is around 2 mV for pulses at 150% RMT at a low background activation rate (Ziemann et al., 1996; Hallett, 2007; Devanne et al., 1997). The shape of the MUAP is described by a first order Hermite-Rodriguez function *H*(*t*) (Li et al., 2012; Moezzi et al., 2017; Olmo et al., 2000):

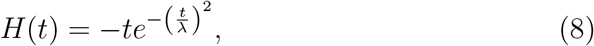

where *λ* is a constant timescale which we set at 2.0 ms (Li et al., 2012). Thus, an electromyogram (EMG) response, *M* (*t*), is given by the sum of the contributions from the various MUAPs:

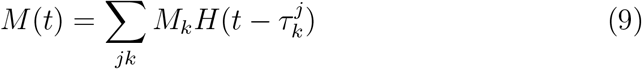

We define a MEP as the maximum positive deflection plus the maximum negative deflection.

We mostly used standard parameter values to match Li et al. (2012), but have adjusted slightly the MUAP amplitude *M*_0_ and the minimum threshold *T*_*min*_ to give very limited EMG activity with no voluntary contraction. To investigate sensitivity of the model to its parameters, we change model parameters by 15% upwards and downwards and analyse the effects on the motor responses.

### 2.4. Underlying assumptions

Before discussing the specific applications of the model, we briefly highlight the most significant underlying assumptions. First, the model is constructed using the I-wave hypothesis of Rusu et al. (2014) for the topology of the connections between layer 2/3 neuronal populations and layer 5 corticospinal output neurons. Secondly, we have used a neural-field approach with the NFTsim model Sanz-Leon et al. (2017), and assume that this is appropriate for modeling TMS. Its applicability to many scenarios has been well discussed in Deco et al. (2008) and Pinotsis et al. (2014), for example for modeling EEG and fMRI responses. Third, we assume a reduced external activation of the layer 5 cells compared with the layer 2/3 cells. Finally, we assume that inhibitory layer 2/3 cells are preferentially stimulated at lower TMS intensities, while the excitatory layer 2/3 cells are more excited at higher intensities. We assess the impact of the final two assumptions by changing the parameter values. In principle, we can also change the topology of the connections, although we do not do this in this work.

### 2.5. Application to single- and paired-pulse protocols

To evaluate how well our model captures TMS-evoked activation of the corticomotor system, we assessed the model’s capacity to generate TMS-related phenomena. First we assessed how the modeled MEP changed with increasing TMS intensities (i.e. an input-output curve). Typically, MEPs increase in sigmoid shape, reaching a plateau above ~180% resting motor threshold (RMT). From the response curve, we identify RMT as the intensity *ϕ*_*x*_ requried to produce a MEP of 0.1 mV in size. While RMT is typically identified as the minimum intensity required to evoke at least 5 of 10 MEPs>0.05 mV in amplitude, our model does not include variability, meaning MEPs are identical in amplitude across simulations provided the parameters are kept constant. Our decision to use 0.1 mV as RMT comes from studies assessing input-output curves which typically observe a mean MEP amplitude of this size at 100%RMT [for an example see Rogasch et al. (2009)]. We also assessed the capacity of the model to predict unseen experimental data using input-output curves from Goldsworthy et al. (2016) (mean MEP amplitudes between 90% RMT to 180% RMT in 10% increments). Specifically, we adjusted the excitatory-to-excitatory and excitatory-to-inhibitory coupling strengths, *ν*_*ee*_ and *ν*_*ie*_ respectively, in order to best fit the experimental data over the range 90% RMT to 130% RMT, using the sum of squares of the deviation of the model from the experimental data (with each point weighted by the inverse square of the standard error in the experimental data) as our measure of goodness of fit. That is, we found *ν*_*ee*_ and *ν*_*ie*_ that minimized

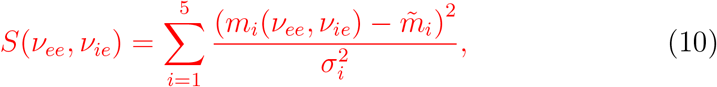

where *m*_*i*_(*ν*_*ee*_, *ν*_*ie*_) is the modeled MEP for the *i*-th intensity (70%, 80%, 90%, 100%, 110% RMT) for couplings *ν*_*ee*_ and *ν*_*ie*_, 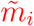 is the mean measured MEP at the *i*-th intensity, and *σ*_*i*_ is the standard error in the mean measured MEP at the *i*-th intensity. To find the approximate position of the minimum of *S*, the parameters *ν*_*ee*_ and *ν*_*ie*_ were varied between 0.75 and 1.05 times their standard values of Appendix A in steps of 0.05 times their standard value, then to refine the position, the parameters were varied in finer steps of 0.01 times their standard value in the vicinity of the minimum. We then used the fitted *ν*_*ee*_ and *ν*_*ie*_ to model MEP amplitudes between 140% RMT and 180% RMT and compared to experimental data. That is, we have predicted data at high intensities that is unseen by the model.

Second, we assessed how the MEP was modulated following paired pulse paradigms. In the paired pulse approach, a sub- or suprathreshold conditioning TMS pulse was delivered followed by a test TMS pulse after a given inter-stimulus interval. The peak-to-peak amplitude of the conditioned MEP is then compared against a MEP following a test TMS pulse alone (i.e. an unconditioned MEP). Experimentally, subthreshold conditioning pulses are followed by a period of inhibition lasting 1-6 ms (short-interval intracortical inhibition, SICI) and then a period of facilitation lasting 10–15 ms (intra-cortical facilitation, ICF) (Kujirai et al., 1993). Suprathreshold conditioning pulses are followed by strong facilitation which peaks at ~20 ms, and then a long period of inhibition lasting 50–200 ms (long-interval intracortical inhibition, LICI) (Valls-Solé et al., 1992). SICI and ICF are increased and decreased respectively by drugs which are agonists of GABA_A_receptors (i.e. increase inhibitory neurotransmission) such as benzodiazepines (Ziemann et al., 1996), and antagonists of *N*-Methyl-D-aspartic acid (NMDA) receptors (i.e. decrease excitatory neurotransmission) such as dextromethorphan (Ziemann et al., 1998). In contrast, LICI is increased by drugs which are agonists of GABA_B_ receptors, such as baclofen (McDonnell et al., 2006).

To test SICI and ICF, we used a conditioning subthreshold pulse at 70% RMT by setting *ϕ*_*x*_ to 70% of its threshold value identified from the response curve, followed by a second, test pulse, at 120% RMT, at a variety of intervals. The MEP due to the second pulse was then plotted against the interstimulus interval. To model the impact of NMDA-modulating drugs on SICI and ICF, we reduced the coupling constant *v*_*ee*_ by 30% and analysed the changes in MEP against ISI. To assess the effect of GABA_A_modulation, we increased the coupling constant 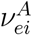 by 30% and assessed the change in MEP. To quantify the effect on SICI, we looked at the reduction in amplitude at 3 ms ISI; for ICF we looked at the change in amplitude at 15 ms ISI.

LICI was analysed in a similar manner. In this case we used a 120%RMT conditioning pulse and a 120% test pulse. The MEP was plotted against ISI. To look at the effect of modulating GABA_B_, we increased the coupling constant 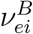 by 30% and simulated the MEP against ISI plot again. We then used the change in amplitude at 100 ms ISI as a measure of the effect on LICI. We note the choice to alter parameters by 30% was not intended to specifically model the effect of these drugs at any specific dose based on biophysical grounds, but was instead selected to generally assess the effect of increasing or decreasing parameters related to the primary mechanism of action for each drug.

### 2.6. Muscle contraction

Finally, we assessed how the modeled MEP was altered with a tonic muscle contraction. MEPs increase in amplitude with increasing voluntary muscle activation, and are followed by a cessation in muscle activity which last for 200–300 ms and is known as the cortical silent period (CSP). The firing rate of the Layer 2/3 cells is directly related to the level of voluntary muscle contraction (Evarts, 1968), and this influences the rates of the Layer 5 cells. Moezzi et al. (2017) used a Poisson process to provide excitatory input to layer 2/3 cells with a mean rate of 5 s^−1^ to model 10 percent maximum voluntary contraction (%MVC). This rate gave similar motoneuron activity to that observed experimientally (Zhou and Rymer, 2004). In our neural field implementation, we have applied an additional external excitatory input rate to the layer 2/3 populations of a constant 0.5 s^−1^ for each %MVC, coupled to the *e* and *i* populations with strengths *ν*_*ee*_ and *ν*_*ie*_ respectively. For example, for 5% MVC, we apply, in addition to the rate provided by the TMS stimulation, a constant rate of 2.5 s^−1^; for 10% MVC the additional rate is 5.0 s^−1^.

### 2.7. Application to repetitive TMS protocols

A central motivation for developing a MEP model of TMS was to provide a more realistic output measure for neural field models of plasticity induced by repetitive TMS (rTMS) protocols. As a proof-of-concept, we assessed how modeled MEPs were altered following either intermittent or continuous theta burst stimulation (iTBS, cTBS). We included features of calcium-dependent plasticity with a Bienenstock-Cooper-Munro (BCM) rule for metaplasticity (Fung and Robinson, 2014; Wilson et al., 2016, 2018), see Appendix B. We and others have previously demonstrated that this plasticity model captures several key features of TBS-induced plasticity by assessing changes in synaptic weights following stimulation (e.g. synaptic weights are increased following iTBS and decreased following cTBS). However, it remains unclear how these changes in synaptic weights would impact MEP amplitude.

We simulated canonical cTBS and iTBS protocols (Huang et al., 2005) with three pulses per burst at 50 Hz intraburst rate, five bursts per second, for a total of 600 pulses. For cTBS pulses were applied continuously; for iTBS pulses were applied for 2 s then were absent for 8 s, before repeating. Specifically, we first measured the ‘pre-TMS’ response of the model to a single test pulse at 120% RMT, that is, a single pulse of *ϕ*_*x*_ with an amplitude of 120% of the value identified from the modeled motor response. We then applied TBS by pulsing the external stimulation rate *ϕ*_*x*_ in a cTBS or iTBS pattern with an amplitude of 80% of the threshold value, that is 80% RMT. Plasticity induced by TBS was modeled using the approach outlined in Wilson et al. (2016), where changes in *ν*_*ee*_ are governed by intracellular calcium concentrations following stimulation (see Appendix B for details). We then identified from NFTsim the final value of 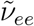, the ultimate synaptic weight for the excitatory-to-excitatory coupling arising as a result of the TBS application. We then used this value as a new value for *ν*_*ee*_, and simulated again the MEP response to a single pulse at intensity of 120% RMT. The test responses before and after TBS were then compared.

Although initial experimental studies suggested that iTBS increased, whereas cTBS decreased, MEP amplitude, more recent studies have suggested that response to TBS is variable across individuals. The variability likely arises from both methodological and biological factors. For instance, Hamada et al. (2013) found that the manner of interaction of TMS with cortical circuits was associated with the direction of change in MEPs following iTBS and cTBS. In contrast, Mori et al. (2011) found that single nucleotide polymorphisms in genes associated with glutamatergic NMDA receptors impacted iTBS outcomes. To assess the impact of methodological and biological variability on TBS-induced changes in MEPs, we altered the synaptic coupling of TMS to the layer 2/3 excitatory population, *ν*_*ex*_, to mimic variability in how TMS interacts with cortical circuits during TBS, and the initial value of the coupling between layer 2/3 excitatory populations, *ν*_*ee*_, to mimic variability in glutamatergic receptors. The change in MEP following TBS was then assessed as a function of these two coupling parameters.

### 2.8. Code availability

The NFTsim model for the neural field equations is available at https://github.com/BrainDynamicsUSYD/nftsim (version 1.1.0) (Sanz-Leon et al., 2017). The motor model is implemented in Matlab and is available at https://github.com/mtwilson1970/MEP_modeling_2020.

## 3. Results

### 3.1. Motor evoked potential at rest

A simulated EMG at rest is shown in Fig. 3 by the black lines. A stimulation intensity of 780 s^−1^ (120% RMT) has been used with a pulse length of 0.5 ms (Wilson et al., 2016). Broadly, these parameters result in activation of around 39% of neurons, of similar magnitude to the fraction of neurons reported to respond to single TMS in non-human primates (Romero et al., 2019). Part (a) gives a plot of the EMG as a function of time (stimulation is at 0 s); the MEP is indicated. The corresponding layer 5 pulse rate is shown in part (b), and the firings of motoneurons are shown in part (c). Each dot corresponds to a firing of a unit arranged such that the lowest threshold firings correspond to the lowest-indexed units. The MEP demonstrates a realistic shape, with a rapid positive rise about 25 ms after the TMS pulse, followed by rapid fall to a maximum negative deflection about 10 ms later. Small amplitude (~0.05 mV) activity follows the MEP, from about 50 to 75 ms after the TMS pulse. In this case the MEP shows a double positive peak, with the main peak at about 1.05 mV and the secondary peak about 0.50 mV in size. While MEPs are typically biphasic in shape, more complex triphasic waveforms are often recorded [for example see Day et al. (1989)].

**Figure 3:**
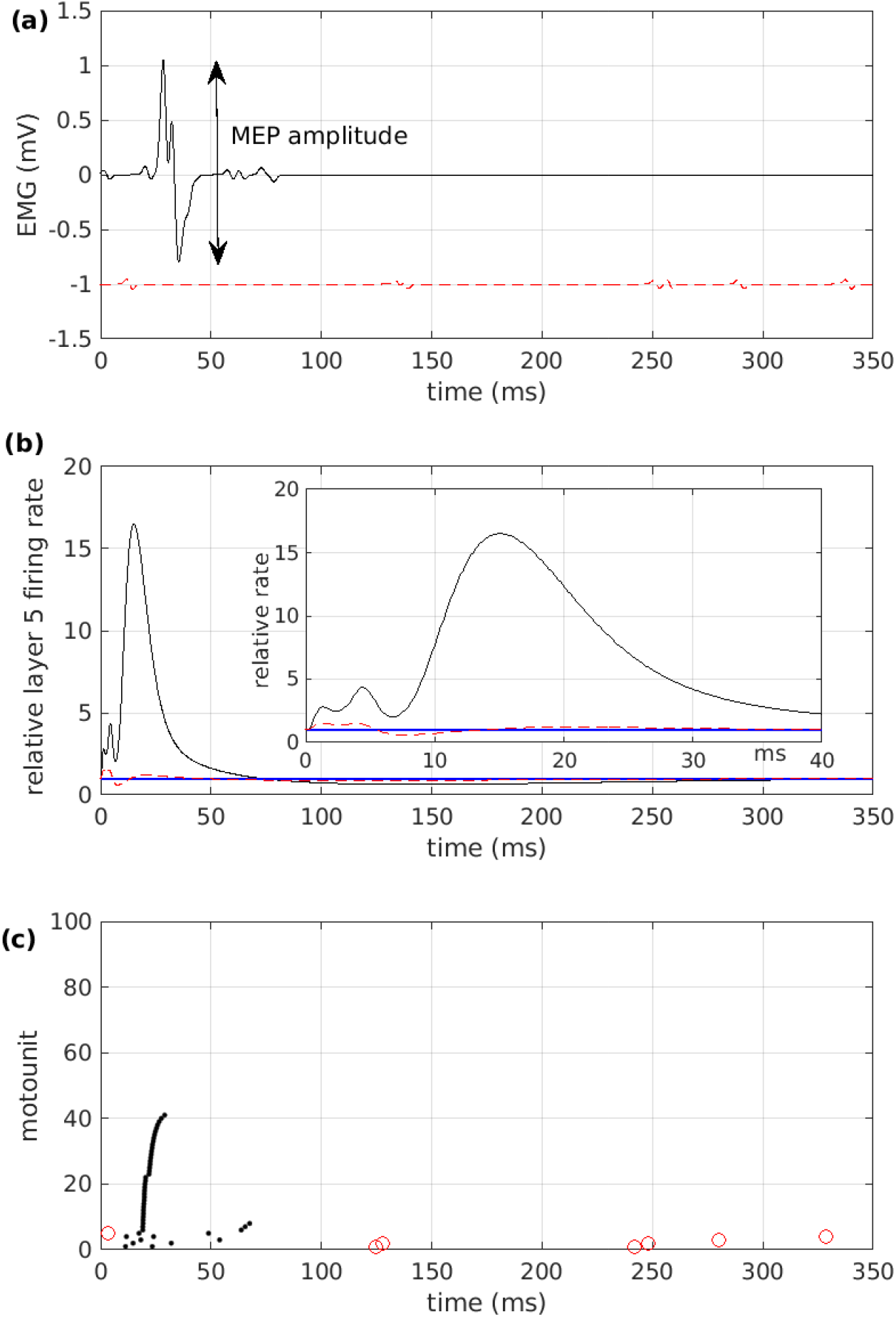
(a) A typical electromyogram (EMG) produced by the model as a function of time, for the suprathreshold case of 120% resting motor threshold (RMT), black line, and the subthreshold case of 80%RMT, red dashed line, displaced by −1 mV. The motor-evoked potential (MEP) is indicated. The t1ra9nscranial magnetic stimulation (TMS) pulse occurs at 0 s. (b) The mean rate of firing of the layer 5 population as a function of time, plotted relative to the baseline rate, for the cases of 120%RMT (black curve) and 80%RMT (red dashed curve). The blue line shows the baseline rate. The inset shows an enlarged version at early times. (c) Motoneuron firings as a function of time, for the cases of 120%RMT (black dots) and 80%RMT (open red circles).

Figure 3(b) shows the mean pulse rate, relative to the baseline of 17.8 s^−1^. Initially, around 2 ms after the initial TMS pulse, there is a small (relative change 2.5) peak in the response. This time delay for layer 5 response compares reasonably with the 2 to 3 ms of Moezzi et al. (2017) and is similar to experiment (1.5–2 ms) (Hallett, 2007). There are further peaks of activity. A peak of 4.5 occurs in relative activity at 5 ms after stimulation, and a peak of nearly 17 at 15 ms. Examination of the modeled cortical responses shows that these peaks are a result of indirect stimulation of the layer 5 cells, via the excitatory layer 2/3 cells. Their size is influenced by the magnitude of the direct stimulation to the layer 5 neurons. There is a strong dip before the final peak, at ~8 ms, due to build-up of GABA_A_ inhibitory effects after the initial rise, but it is short-lived. After about 15 ms, the layer 5 neurons reach a maximum firing rate. Beyond this time, there is a drop-off in activity as the longer timescale GABA_B_ inhibitory receptor effects become prevalent. Since Fig. 3(b) shows mean activity across a population, the peaks and troughs of the plot should not be considered as the I-waves *per se*; rather we may expect such waves to be possible during the times where the activity is large. Changes in layer 5 firing rate are consistent with recent invasive recordings in rodents (Li et al., 2017) and non-human primates (Romero et al., 2019).

The firings of the motoneurons are shown in Fig. 3(c). Note that the motoneurons fire in approximate synchrony at the MEP, although the higher-threshold neurons (higher motounit number) fire slightly later, since they need to integrate more input before they reach their thresholds. This is the reason for the ‘double peak’ in the MEP in Fig. 3(a); the later firing neurons contribute larger MUAPs and so the net summation of the MUAP train has a second peak at later times.

Fig. 3 also shows a result for the case of a subthreshold pulse at 80%RMT (red dashed line). In this case we see that the layer 5 firing rate increases by about 50% for the first 6 ms and then decreases by about 50% from baseline for the next 4 ms, panel (b). However, it is insufficient to substantial firing of motoneurons (c) and thus no MEP is produced (a).

The effect of TMS intensity on MEP amplitude is shown in Fig. 4(a). In this plots we show the MEP amplitude against external stimulation rate *ϕ*_*x*_ for a variety of parameter sets. To prevent a single plot becoming exceptionally crowded, we shown just a small subset of the parameter sets. The full analysis is shown in Appendix C. The standard parameter set (see Appendix A) produces a response curve shown by the thick black line. From this curve we identify a resting motothreshold of about 650 s^−1^ input intensity, as being the amplitude that gives a MEP of around 0.1 mV. We show with vertical dashed lines this value and 120% of this value.

**Figure 4:**
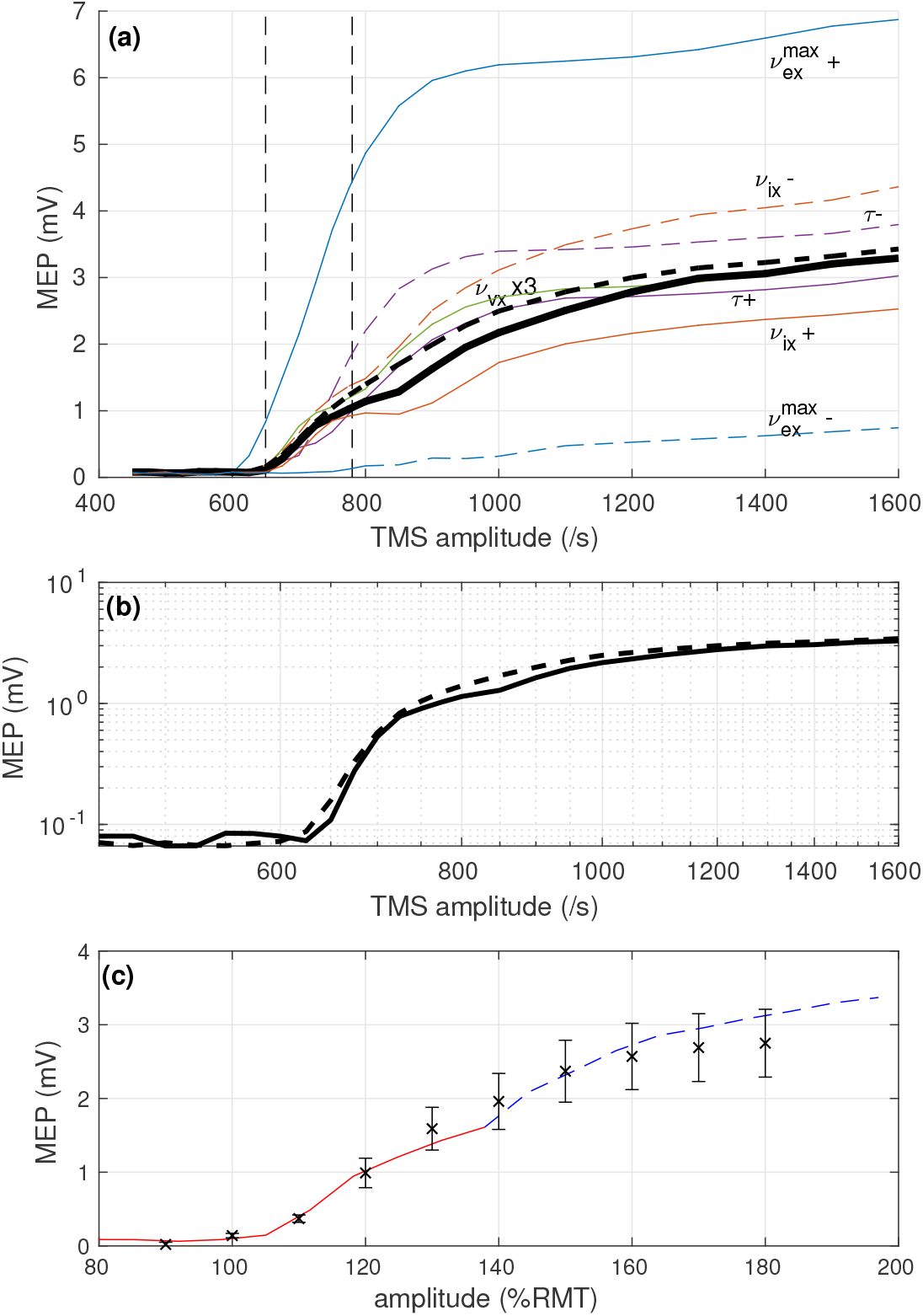
(a) The size of the modeled motor-evoked potential (MEP) as a function of the stimulation intensity at rest, for various parameter sets. The black solid line denotes the response for the standard parameter set. Shown by the colored lines are the response curves due to: a +15% change in maximum external to excitatory coupling 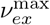, blue solid, labeled 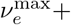; a −15% change in 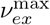, blue dashed, labeled 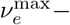; a +15% change in external to inhibitory coupling *ν*_*ix*_, orange solid, labeled *ν*_*ix*_+; a −15% change in *ν*_*ix*_, orange dashed *ν*_*ix*_−; setting layer 2/3 to layer 5 axonal delays 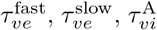 and 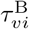 all to 3 ms, purple solid, labeled *τ* +; setting 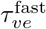 to 1 ms, 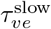 to 6 ms, and leaving 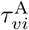 and 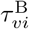 at 3 ms, purple dashed, labeled *τ* −; multiplying external to layer 5 coupling *ν*_*vx*_ by three, green solid, labeled *ν*_*vx*_ × 3. The black dashed line denotes the mean response across all parameter sets modeled. The vertical dashed lines indicate resting motor threshold (RMT) and 120%RMT for the standard parameter set. (b) The same plot as part (a) on a log-log scale for the standard parameter set (solid line) and the mean across parameter sets (dashed line). (c) A prediction of high amplitude single-pulse response based on the low amplitude single-pulse response. The b2la1ck points show experimental data (mean standard error in the mean) for a single-pulse response curve as measured by Goldsworthy et al. (2016). The red line shows model output after varying the synaptic couplings *ν*_*ee*_ and *ν*_*ie*_ to best fit the low amplitude experimental data points (up to 130% RMT), based on a least-squares measure. The model then reproduces the experimental measurements at the higher amplitudes (140% RMT and above), shown by the blue dashed line, within experimental uncertainty.

The colored curves show the response curves when changes are made to the parameter set. Except for the propagation delays 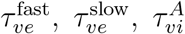 and 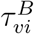, we have changed one parameter at a time, either increasing it by 15% or decreasing it by 15%. With the propagation delays we have changed them by setting (i) all equal to 3 ms and (ii) decreasing 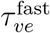 to 0 ms and increasing 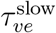 to 6 ms (i.e. (i) equalizing them and (ii) stretching them). We also show the effect of increasing *ν*_*vx*_, the TMS-to-corticospinal output neuron coupling, by a factor 3. The black dashed line shows the mean over the responsee curves for all parameter sets. A more complete analysis is given in Appendix C.

There is considerable variation in maximum output intensity, from about 1 – 7 mV. However, there is more consistency in threshold; all responses are very low at 600 /s and then climb rapidly with stimulation intensity. By 1200 s^−1^ stimulation (approx 180% RMT) most responses have flattened. For clarity, Fig 4(b) shows the MEP response for the standard parameter values and mean of the responses for the ±15% changes in parameters, that is the thick black and dashed lines of part (a), on a log-log scale.

Appendix C shows a more complete sensitivity analysis for changes in model parameters. Response curves are shown in Fig. C.9. In Table C.3, we document the percentage changes in 1. resting motor threshold (RMT), 2. MEP amplitude at 200% threshold and 3. maximum gradient of the response curve for the various changes in parameters. The RMTs are not greatly affected by changes in parameters, with *θ*_*v*_, the ‘threshold’ of the layer 5 activation curve being the most significant contributer. The MEP amplitude at 200% RMT, (towards the ‘plateau’), shows greater variation. In particular, parameters associate with the balance between excitatory and inhibitory coupling in layer 2 and 3 affect the MEP amplitude, a set of parameters favouring excitation gives a strong increase in MEP, and the opposite with inhibition. Also, the strength of the TMS-to-excitatory coupling, 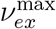, strongly affects MEP size (the greater the field the strength, the greater the response), as does the threshold of the layer 5 activation curve, *θ*_*v*_. The maximum gradient in response curve can be strongly affected by choice of parameters, in particular the strength of the TMS-to-layer 2/3 excitatory coupling 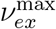, the time delays for the layer 2/3 excitatory to layer 5 couplings and the threshold of the layer 5 activation curve, *θ*_*v*_. This manifests itself on Fig. 4(a) as a change in the rate of rise of MEP with increasing stimulus above threshold. This would strongly affect the rate of increase of MEP just above threshold and thus the extent of stimulation at 120% RMT.

Finally, we tested the predictive capacity of our model against experimental input-output curve data taken from Goldsworthy et al. (2016). The optimal fitting of MEP amplitudes using data obtained between 90% and 130% RMT gave *ν*_*ee*_ and *ν*_*ie*_ as 0.92 and 0.89 times the standard parameter values of Appendix A respectively. With these values, we are able to predict the experimental response for the higher stimulation amplitudes, 140% RMT to 180% RMT, that is, we have predicted the response to the higher intensities based on the response to the lower intensities. Figure 4(c) shows the results; the black points show the experimental data (mean ± standard error in the mean), the red curve shows the model output fitted to the lower amplitude data (90% RMT to 130% RMT), and the blue dashed continuation of the red curve gives the prediction for the high amplitude data (140% RMT to 180% RMT). The prediction lies within experimental uncertainty, providing evidence that the model is capable of predicting unseen data.

### 3.2. Paired-pulse protocols

To model SICI and ICF we have applied a conditioning stimulus at 70% RMT and a test stimulus at 120% RMT. The interstimulus interval (ISI) has been varied up to 20 ms. Results are shown in Fig. 5(a). For short ISI (5 ms or less) there is substantial inhibition of the pulse; at very short ISI the response to the test pulse is almost abolished. This broadly agrees with experiment (shown for comparison) which demonstrates that SICI at subthreshold conditioning intensities persists up to approximately 7 ms ISI (Kujirai et al., 1993). At longer ISI (7 ms and greater) the model shows facilitation of the test pulse. This facilitation peaks at about 10 ms, in agreement with experiment (Valls-Solé et al., 1992; Kujirai et al., 1993; Ziemann et al., 1996).

**Figure 5:**
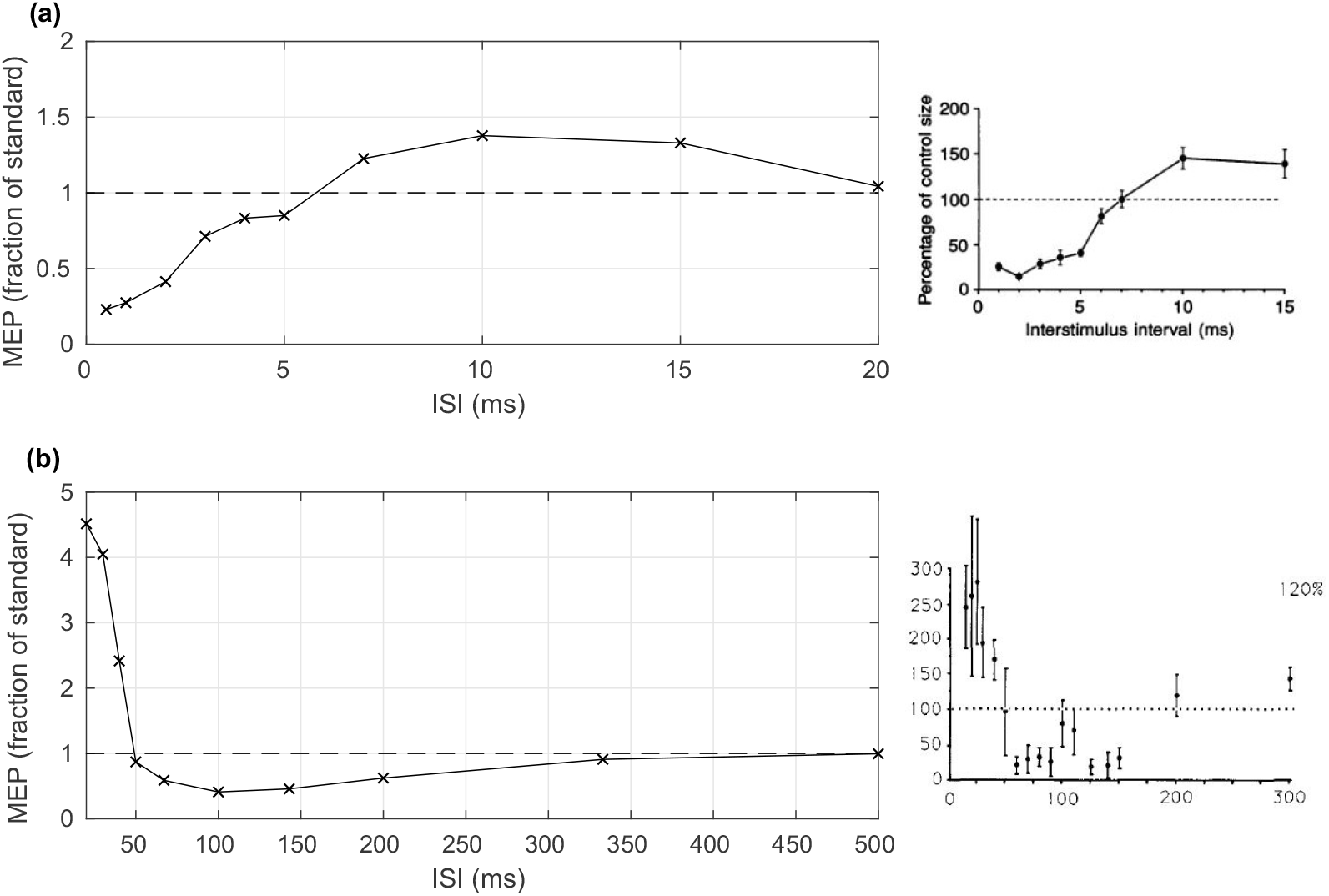
The modeled motor-evoked potential (MEP) changes as a result of paired-pulse protocols. MEPs are normalized in terms of the amplitude for a single test pulse and are plotted against the time between the two pulses, that is the interstimulus interval (ISI). (a) A 70% resting motor threshold (RMT) conditioning pulse with a 120% test pulse demonstrates short interval intracortical inhibition (SICI) and intracortical facilitation (ICF). Experimental results (Kujirai et al., 1993) are shown in the right hand panel with permission. (b) A 120% RMT conditioning pulse with a 120% RMT test pulse shows long interval intracortical inhibition (LICI). The dashed lines indicate no change in MEP; a response above the line indicates facilitation, a response below indicated inhibition. Experimental results (Valls-Solé et al., 1992) are shown in the right hand panel with permission.

Modeling of LICI is achieved by pairing two suprathreshold pulses at 120% RMT. Results are shown in Fig. 5(b) and an experimental plot (Valls-Solé et al., 1992) is shown for comparison. At ISI less than 50 ms there is substantial facilitation of the test pulse, but at longer ISI (50 ms to 300 ms) there is considerable inhibition. While broadly consistent with experiment there are some differences. First, the extent of the ICF is higher than usually seen, and the period of LICI lasts to longer ISI (about 300 ms) than is typically seen experimentally (about 200 ms). Also, the modeled LICI is not as strong; MEPs are reduced in the model to around 40% of their baseline whereas in experiment they can be almost eliminated.

Next, we simulated the effect of GABAergic and anti-glutamatergic drugs on SICI, ICF, and LICI by modulating the coupling strengths between layer 2/3 cortical populations. Fig. 6(a) shows a paired pulse response curve with and without the presence of a GABA_A_ agonist, modeled by increasing the coupling strength 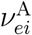 due to GABA_A_ receptors from the inhibitory to the excitatory population. The plot shows that increasing neurotransmission of GABA_A_ receptors on excitatory populations increases SICI (at 3 ms) but reduces ICF (at 15 ms). This agrees with experiment (Ziemann et al., 2015) which shows that a GABA_A_ receptor agonist such as Diazepam increased SICI (Di Lazzaro et al., 2005, 2007; Müller-Dahlhaus et al., 2008) but reduced ICF (Inghilleri et al., 1996; Mohammadi et al., 2006).

**Figure 6:**
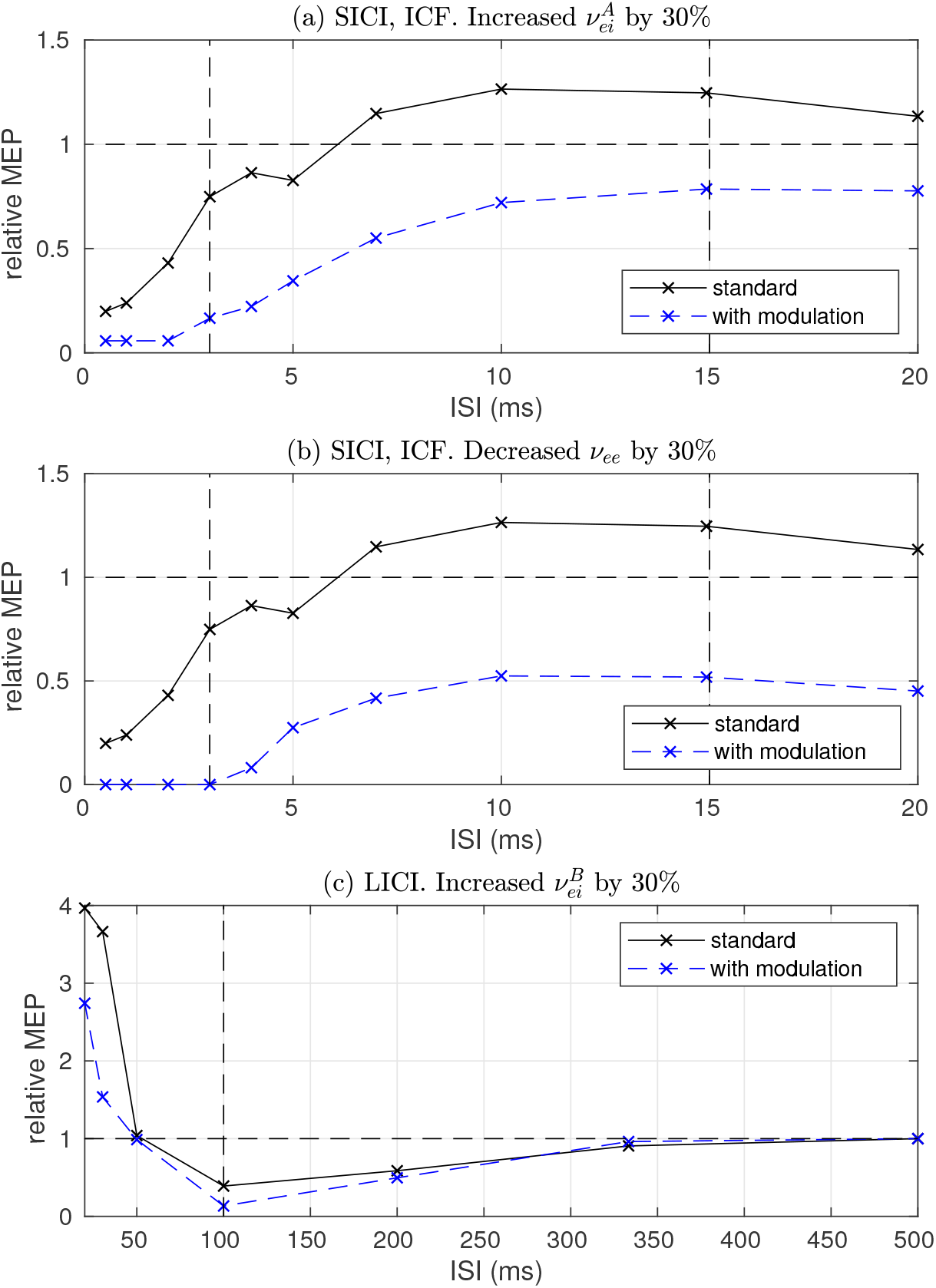
Paired pulse response curves with and without the presence of modulating drugs. (a) The relative motor evoked potential (MEP) as a function of interstimulus interval (ISI) for a subthreshold conditioning pulse and suprathreshold test pulse for the case of the standard parameter set (black solid) and increase of GABA_A_-modulated inhibitory to excitatory coupling 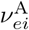 by 30% (blue dashed). The vertical dashed lines indicate ISI of 3 ms (for short-interval intracortical inhibition, SICI) and 15 ms (for intracortical facilitation, ICF). The horizontal dashed line shows a relative MEP size of 1 (no change from baseline). (b) The paired-pulse response for the standard parameter set (black solid) and decrease of excitatory to excitatory coupling *ν*_*ee*_ by 30% (blue dashed). (c) The paired-pulse response for two suprathreshold pulse2s5for the standard parameter set (black) and increase of GABA_B_-modulated inhibitory to excitatory coupling 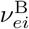 by 30% (blue). The vertical dashed line denotes an ISI of 100 ms (long-interval intracortical inhibition, LICI).

Fig. 6(b) shows the effect of decreasing excitatory to excitatory coupling strength *ν*_*ee*_ (equivalent to applying an anti-glutamatergic drug) on SICI and ICF respectively. The plot shows that SICI (at 3 ms) is increased, whereas ICF (at 15 ms) is reduced by decreasing excitatory coupling, largely in agreement with studies applying anti-glutamatergic drugs such as Mematine (Schwenkreis et al., 1999) or Riluzole (Schwenkreis et al., 2000; Liepert et al., 1997), and NMDA-antagonists such as Amantadine (Reis et al., 2006) or Dextromethorphan.

Finally, we simulated the effect of increasing the inhibitory to excitatory coupling 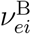 due to GABA_B_ receptors (equivalent to applying a GABA_B_ receptor agonist) on LICI. Figure 6(c) demonstrates a significant increase in LICI (ISI of 100 ms; two suprathreshold pulses) with increased GABA_B_ receptor coupling to excitatory populations, again in agreement with experiments using GABA_B_ receptor agonists such as baclofen (McDonnell et al., 2006). Taken together, these findings demonstrate that our MEP model is able to capture a large range of paired-pulse TMS phenomena that are observed experimentally, including the effects of altering excitatory and inhibitory neurotransmission using different drugs.

### 3.3. Motor evoked potential during contraction

Figure 7(a) show the EMG response following a single pulse at 120% RMT during a voluntary tonic muscle contraction (10% MVC). Before the pulse, ongoing muscle activity of around 0.05 mV in amplitude is apparent, as a result of the excitatory input to the cortex from the voluntary contraction (modeled as an external input rate of 5 s^−1^ on the excitatory cortical population.) The background activity has resulted in a 10% increase in the amplitude of the MEP during a contraction. Also, a silent period is evident after the pulse during which there is no EMG, with background EMG returning after 300 ms. Further increasing the strength of the muscle contraction resulted in increased MEP amplitude, as shown in Fig. 7(b), in line with experimental findings.

**Figure 7:**
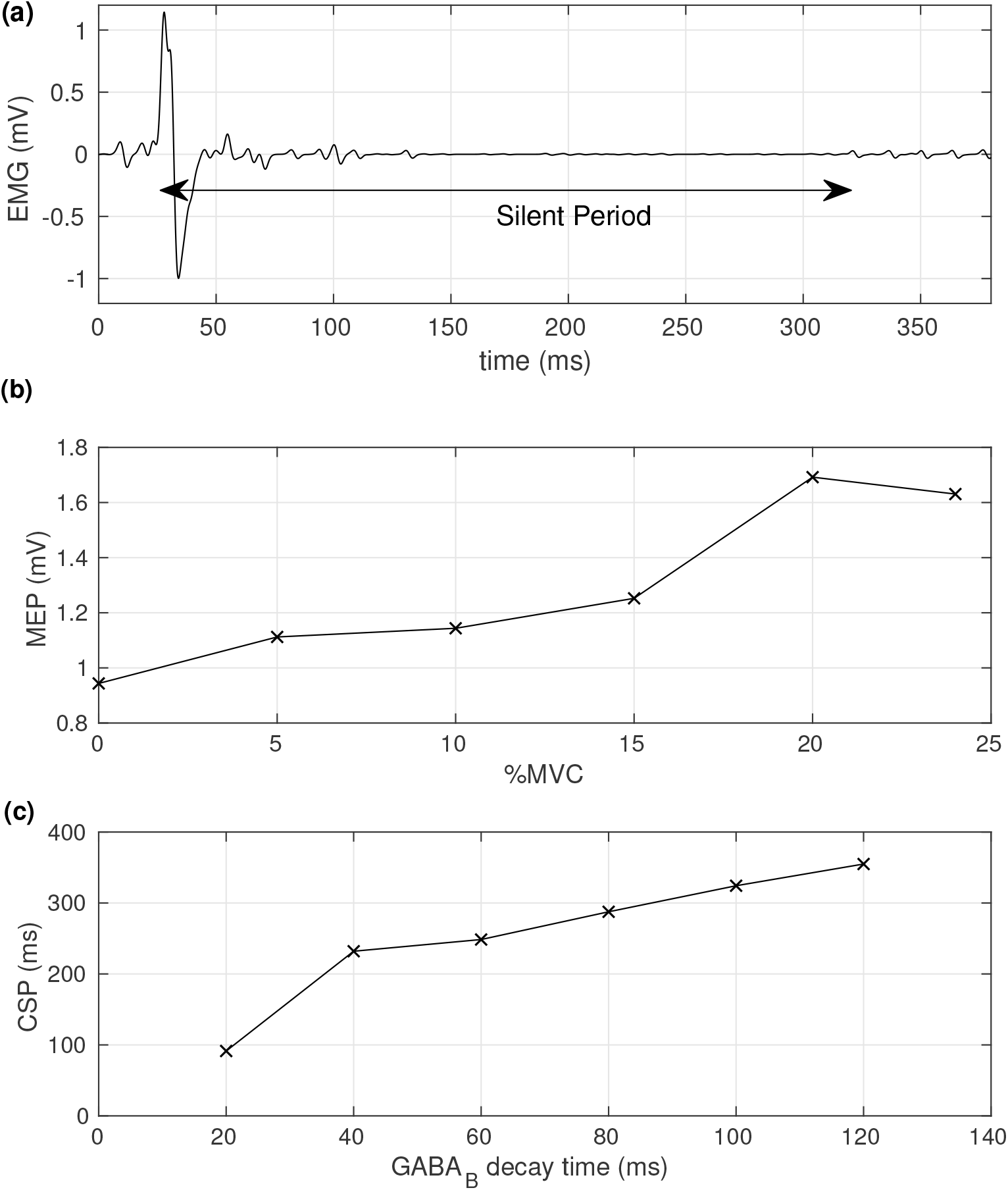
Modeled motor-evoked potential (MEPs) with muscle contraction. (a) The time-course of the electromyogram (EMG) at 120% resting motor threshold (RMT) intensity and 10% maximum voluntary contraction (MVC). The silent period is indicated. (b) The amplitude of the MEP as a function of %MVC. (c) The duration of the silent period as a function of *γ*-aminobutyric acid B (GABA_B_) decay time constant.

Figure 7(c) shows the CSP against the time constant of the decay of GABA_B_. As decay constant increases, the silent period also increases. This agrees with experimental results, but overall the modeled CSP is somewhat longer than typical measured CSPs (Li et al., 2017). The CSP reflects the long period in which the layer 5 pulse rate (Fig. 3(b)) drops below its equilibrium value due to build-up of GABA_B_ and its length is strongly related to the timescale of GABA_B_ decay (Moezzi et al., 2017).

### 3.4. Theta-burst stimulation

Having established that our model captures a wide range of single and paired pulse TMS phenomena, we next assessed whether modeled MEPs at 120% RMT were sensitive to changes in synaptic weights induced following 600 pulses of canonical iTBS and cTBS (Huang et al., 2005) stimulated using a model of CaDP with metaplasticity. We modeled a range of parameter values for *ν*_*ex*_ and *ν*_*ee*_ mimicking variability in how TMS interacts with cortical circuits (TMS-e coupling) and variability in glutamatergic neurotransmission (e-e coupling) respectively. Figure 8 shows the predicted relative changes in MEPs following both cTBS and iTBS. There are several notable features to these outcomes. First, there are several areas within the parameter space that predict the ‘canonical response pattern to TBS (i.e. cTBS decreases MEPs, iTBS increases MEPs, e.g. the point). Second, a wide variability in response profile can also be generated by altering how TMS interacts with cortical circuits (TMS-e coupling) and how excitatory populations interact with each other (e-e coupling). For instance, the point * shows an opposite to canonical response, the point ∇ a parameter set where both paradigms decrease MEP amplitude, and ∆ a parameter set where both paradigms increase MEPs. Furthermore, there are parameter spaces where neither paradigm has a strong effect on MEP amplitude (e.g. ‘non-responders’). However, the maximum predicted increase in MEPs (~1.15) is smaller in magnitude than the maximum predicted decrease (~0.7), and is also smaller than the maximum often observed in experiment (~1.8). Third, the response of NFTsim to TBS becomes unstable (i.e. locks in to a high firing rate similar to a seizure) at high values of both TMS-e coupling, but particularly e-e coupling, due to the intracortical excitatory-to-excitatory feedback overcoming the inhibitory-to-excitatory feedback (Roberts and Robinson, 2012; Robinson et al., 2004). This stiuation is shown by the white space on Fig. 8 since for these cases we cannot assess changes in MEPs after TBS. It occurs when potentiation is strong, and patly explains why the maximum MEP increase shown in the model (~1.15) is rather lower than often seen experimentally (~1.8). Interestingly, disorders associated with abnormal glutamatergic receptor function, such as anti-NMDA receptor encephalitis, are often accompanied by seizures. Fourth, the predicted changes in MEP amplitude following TBS across the parameter space are nonlinear — suggesting these relationships would not be evident with simple correlations often used in human TMS experiments. Taken together, these findings demonstrate that our MEP model is sensitive to changes in synaptic weight following TBS predicted by a model including rules for CaDP and metaplasticity, and is able to capture a variety of response profiles often observed in human TBS experiments. However, the model is not able to adequately capture the magnitude of MEP changes observed in experiments following TBS, suggesting that modeling plasticity only on excitatory-to-excitatory synapses is likely insufficient to explain experimental observations.

**Figure 8:**
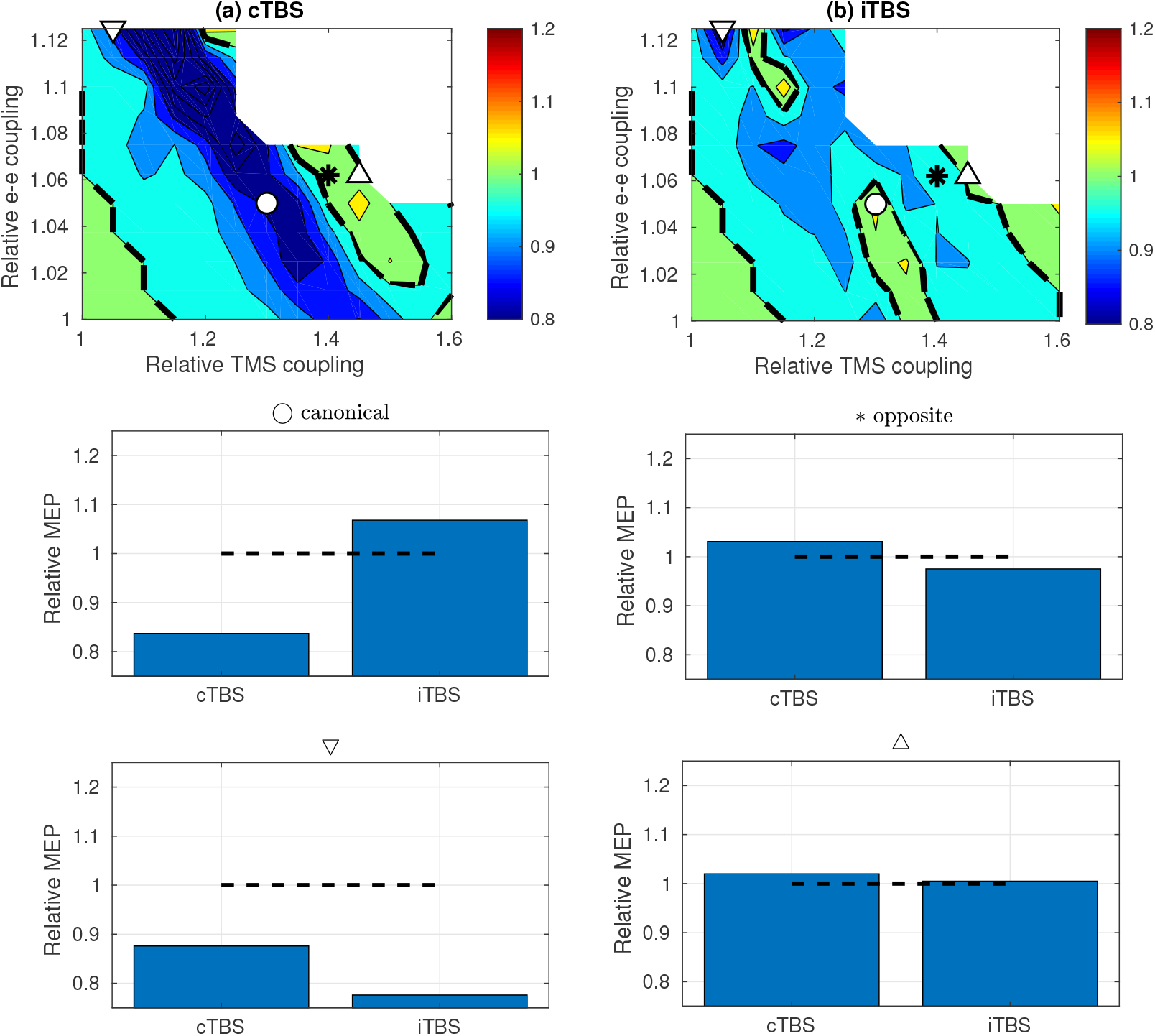
The modeled MEP changes as a result of (a) continuous theta-burst stimulation (cTBS) and (b) intermittent theta-burst stimulation (iTBS) protocols against the relative strength of the transcranial magnetic stimulation (TMS)-to-excitatory coupling and excitatory-to-excitatory coupling strengths. The dashed lines show no change in synaptic weight. The weight changes at four points are indicated in the lower panels. A canonical response is indicated by ◯, an opposite-to-canonical response is indicated by *, both responses positive is indicated by ∆, and both responses negative by ∇. The white space in panels (a) and (b) denotes regions where the output from the NFTsim simulation locks into a persistent high-firing state and MEP changes cannot be assessed.

## 4. Discussion

We have developed a biophysical model of MEPs following TMS to the motor cortex by combining a population-based model of cortical activity and an individual neuron model of motor output. The model captures many common features of MEPs including input-output characteristics, responses to paired-pulse paradigms, a silent period with voluntary contraction and changes in MEPs following plasticity-inducing TBS paradigms. It provides unique insights into how micro/mesoscale mechanisms, such as differences in synaptic weightings between excitatory/inhibitory neural populations, can impact TMS-evoked motor output and TMS-induced plasticity.

There are several limitations of, and assumptions behind, the approach. First, we have assumed the architecture of connections between layer 2/3 neural populations and layer 5 corticospinal output neurons used in the I-wave hypothesis of Rusu et al. (2014). We note, however, that varying the lengths of the propagation delays used for the various layer 2/3 to layer 5 connections did not greatly affect the MEP amplitude or threshold, although it did affect the gradient of the response (MEP increase per unit increase in TMS amplitude). Within the neural-field NFTsim model, one could also vary the architecture of the couplings between layers if required.

Secondly, coupling between the external stimulation provided by the TMS coil and the layer 2/3 cells, and layer 5 cells, is not completely understood. This manifests itself in three distinct ways in the model. (a) The interpretation of the external stimulation strength, *ϕ*_*x*_ is moot. In our interpretation of Fig. 4, we have assumed that it is broadly proportional to the strength of the TMS machine output. This assumption affects the identification of the threshold and interpretation of what different percentage RMT might mean in practice. (b) We have made several assumptions regarding the recruitment of inhibitory and excitatory neuronal populations with differing stimulation intensities. Specifically, we have assumed that layer 2/3 inhibitory interneurons are preferentially recruited at lower TMS intensities, whereas layer 2/3 excitatory interneurons become preferentially recruited at higher intensities, Eq. (1). (c) Furthermore, we assume that layer 5 pyramidal cells are only weakly activated by the TMS pulse. While there is evidence that the layer 5 cells receive a lower direct stimulation (Di Lazzaro and Rothwell, 2014; Bungert et al., 2016; Opitz et al., 2013) this is poorly understood. The assumptions (b) and (c) are largely based on indirect experimental observations in humans, e.g. (Ziemann et al., 1996; Ilic et al., 2002), as there is little direct experimental data informing how TMS impacts firing rates of different neuronal types. Recently, both rodent (Li et al., 2017) and non-human primate (Romero et al., 2019) models have been developed which allow recording of individual neurons during TMS. Future research addressing how TMS intensity impacts the firing rates of different neuronal subtypes would greatly inform how best to model these interactions. We have assessed the impact of the assumptions by changing the parameters in the equation, as shown in Fig. 4 and Table C.3. While the size of the MEP and its gradient are sensitive to the maximum external-to-layer 2/3 excitatory coupling size 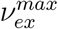, Eq. (1), the other parameters describing the width and threshold of the coupling curve, *B* and *A* respectively, have little impact. There is also moderate impact in the MEP size due to changes in the external-to-layer 2/3 inhibitory coupling, *ν*_*ix*_. Essentially, the stronger the excitation over inhibition, the stronger the MEP. We have assessed the impact of the assumption of low external-to-layer 5 coupling strength by increasing *ν*_*ex*_ by a factor 3; overall MEP size at 120% RMT increased by 20%.

Thirdly, a limitation of our model is that it does not predict I-wave activity in layer 5 corticospinal neurons (Rusu et al., 2014; Moezzi et al., 2017). Population-based models are not well suited for capturing highly synchronized events, such as I-waves, as mean population firing-rates are modeled instead of individual firing events. However, modeling at the population level is well suited for TMS (Wilson et al., 2018), which simultaneously activates large neural populations, and captures the slower excitatory and inhibitory postsynaptic potentials that are likely involved in paired pulse phenomena. Indeed, our model is successful at capturing a wide range of single-pulse, paired-pulse and rTMS phenomena without explicitly modeling I-waves. Furthermore, our model also predicts changes in layer 5 corticospinal neuronal firing rates following single pulse TMS. While it is not currently possible to record this data in humans, the pattern of activity described by the model is remarkably similar to recent in vivo recordings in rodents and non-human primates (Li et al., 2017; Romero et al., 2019). Such findings highlight the potential of such models to inform the microscale and mesoscale mechanisms of TMS from macroscale recordings. We also emphasize that we have used the structure of layer 2/3 neurons feeding forward to layer 5 neurons, as hypothesized by Rusu et al. (2014). However, the essence of our approach, linking neural field theory to MEPs, does not require such a structure, and other structures of connections can be simulated and assessed in a similar manner.

Fourthly, our model is deterministic in the sense that baseline cortical excitability, captured by the mean population firing rate of a given neural population, is relatively constant and has little variability (see the blue line in Fig. 3(b) for example). Thus trial-to-trial variability of MEP amplitude, which is likely driven in part by fluctuations in cortical excitability, has not been considered. Importantly, higher levels of baseline MEP variability are associated with stronger plasticity responses following theta burst stimulation (Hordacre et al., 2017), suggesting neuronal variability plays an important role in promoting brain plasticity. MEP variability is likely driven by fluctuations in both cortical and spinal excitability (Kiers et al., 1993). For example, Zrenner et al. (2018) have demonstrated that the size of MEP varies with the phase of underlying alpha rhythms in the subject, and brain state may influence other TMS effects (Ziemann et al., 2015). In terms of modeling, explicit variations in parameters can be made to investigate the effects of changes in endogenous cortical excitability on MEPs. We have demonstrated that variations in some parameters, e.g. maximum TMS-to-excitatory coupling, 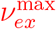, and the threshold of the layer 5 activation curve (*θ*_*v*_; a key parameter determining mean population firing rates), strongly affect MEP sizes and to a lesser extent RMT, suggesting that changes in endogenous cortical excitability has a large impact on MEPs, and variations of these parameters may be important for simulating MEP variability. Furthermore, we also modeled the effect of increased cortical excitability resulting from volitional drive to the motor cortex during voluntary muscle contractions. Increasing the mean population firing rates in the layer 2/3 excitatory population with increasing contraction strength increased MEP amplitude in the model, further demonstrating how changes in endogenous cortical excitability (i.e. brain state) can influence TMS-evoked motor output. Given the importance of variations in cortical excitability for brain function, a necessary next step will involve incorporating variability into the current model. Indeed, such variability may be essential for accurately simulating the brain’s response to TMS. There are multiple ways in which variability in cortical and spinal excitability could be incorporated, such as adapting existing population-based models including cortico-thalamic loops (Robinson et al., 2004), which are able to capture complex fluctuations in brain excitability such as cortical oscillations. Furthermore, more detailed models of spinal circuits could also introduce variability (Kiers et al., 1993). Including these approaches will allow allow further exploration of MEP variability, and modeling of TMS-evoked EEG activity (Rogasch and Fitzgerald, 2013).

Fifthly, we have assumed that the neural population modeling scheme of NFTsim is adequate for the purposes of modeling TMS effects. The relevance of NFTsim to various applications has been well discussed elsewhere (Sanz-Leon et al., 2017; Robinson et al., 2004; Pinotsis et al., 2014; Deco et al., 2008); in particular we note that the application of neural field modeling with calcium dependent plasticity to TMS has been discussed in Fung et al. (2013), Fung and Robinson (2014), and Wilson et al. (2016). We emphasize that we have modeled changes only to the excitatory-to-excitatory coupling *ν*_*ee*_; in practice other coupling strengths are also likely to change (e.g. changes in coupling with inhibitory populations).

Finally, most of the comparisons between model predictions and real data were qualitative in the current work. Our preliminary analysis on the capacity of the model to predict unseen MEP amplitudes resulting from high intensity stimulation when only constrained on low intensity experimental data suggests that the model does have some predictive ability. Furthermore, the model was able to qualitatively capture a wide range of experimental phenomena (input-output curves, paired pulse paradigms at different ISIs and intensities, cortical silent period) using the same parameter set, providing additional support that the model has good predictive capacity. However, a more rigorous and detailed examination of this issue is required. Future work assessing the capacity to predict a wide range of unseen experimental data (e.g. following changes to a large range of experimental parameters such as intensity or inter-stimulus interval) will help further define the predictive value of such models.

## 5. Conclusions

We have demonstrated how a biophysically plausible nonlinear model of MEPs can be combined with the output of a population-based model of cortical neurons in order to produce a description of MEPs due to TMS. The final MEP activity is realistic in terms of variation with intensity and muscle contraction, and demonstrates the known amplitude and interval-dependent effects in paired-pulse stimulation. The MEP model is also sensitive to changes in synaptic weight predicted by a model of TBS-induced plasticity including rules for CaDP and metaplasticity, demonstrating complex relationships between variability in methodological and biological factors and MEP changes following TBS. Overall, the approach allows population-based modeling of cortical plasticity using neural field theory to be better-interpreted, by providing a route by which the effect on the MEP can be evaluated. Continued development of such models in combination with human experiments will enable a unified theoretical understanding of how TMS interacts with and modifies cortical circuits.

## Author statement

MTW constructed the model, carried out most of the modeling, and led the writing of the manuscript. BM provided guidance on the modeling approach. NCR provided experimental background and contributed to the manuscript.

## Conflict of interest

We declare no conflict of interest in this work.

## Acknowledgments

NCR is supported by an Australian Research Council DECRA fellowship (DE180100741). We thank Mitchell Goldsworthy for access to the raw data of Goldsworthy et al. (2016).

## Appendix A. The NFTsim model

The open source NFTsim model, see Sanz-Leon et al. (2017) for details, uses a neural field approach to calculate population-averaged descriptions of neural behavior as a function of time and space. However, in this work we have excluded explicit spatial variation. Here we present briefly the model and its parameters.

The mean soma potential *V*_*a*_ of a population of neurons of type *a* (*a* = *e*, *i*, *v*) for layer 2/3 excitatory, layer 2/3 inhibitory, and layer 5 excitatory corticospinal output neurons respectively, is given by the sum of contributions from postsynaptic potentials (PSPs):

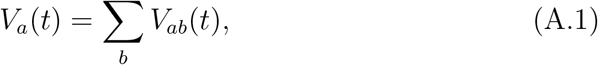

where *V*_*ab*_ describes the postsynaptic potential at the population *a* due to incoming events from population *b* (*b* = *e*, *i*, *v*, or *x*) where *x* describes an external stimulation due to TMS.

The time course of the PSPs is described by the dendritic response through the following equation:

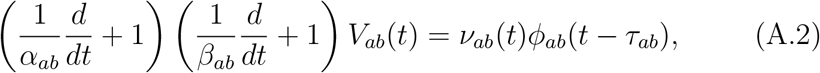

where *α*_*ab*_ and *β*_*ab*_ are rate constants for the rise and fall rates of the PSPs respectively, *ν*_*ab*_ is the strength of coupling, and *ϕ*_*ab*_ describes the rate of incoming axonal pulses, from cells of type *b* to cells of type *a*. The parameter *τ*_*ab*_ describes delay in propagation of signals from neurons of type *a* to type *b*; due to, for example, spatially long nerve pathways.

The mean firing rate of the population *a* is given by a sigmoidal function of the mean soma potential:

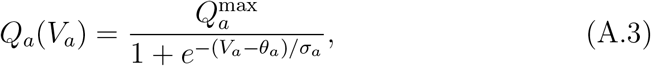

where 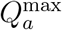 is the maximum firing rate, and *θ*_*a*_ is the threshold of the distribution and *σ*_*a*_ is proportional to the width of the sigmoid distribution.

Finally, the firing of the population *a* results in generation of axonal pulses, which propagate along axons towards synapses. This propagation is described by

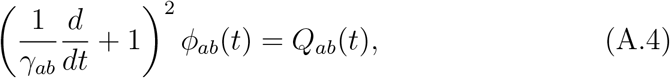

where the parameters *γ*_*ab*_ represent the axonal propagation rates from neurons of type *b* to type *a*. Equations (A.1)–(A.4) describe the time-variation of quantities *V*_*ab*_, *V*_*a*_, *Q*_*a*_ and *ϕ*_*ab*_, and are integrated forward in time to give the behavior of the neural populations.

To simplify, we assume that the dynamics depends only on the presynaptic cell, so that the subscript *a* on the parameters *α*_*ab*_, *β*_*ab*_ and *γ*_*ab*_ is redundant; we label these parameters as *α*_*b*_, *β*_*b*_ and *γ*_*b*_ respectively.

We include a superscript A or B on the synaptic inhibitory parameters to denote the GABA_A_ and GABA_B_ responses respectively. Furthermore, the multiple couplings from the layer 2/3 excitatory cells to the layer 5 cells are labelled with superscripts ‘fast’ and ‘slow’.

The standard parameters for the description of the layer 2 and 3 excitatory and inhibitory cells have come from previous work (Fung et al., 2013; Fung and Robinson, 2014; Wilson et al., 2016, 2014), and are listed in Table A.1. The layer 5 parameters have been chosen to give a plausible motoneuron output, consistent with Moezzi et al. (2017), including a large maximum firing rate, but also with a low threshold for activation. The synaptic couplings to the layer 5 cells from the layer 2/3 cells have been tuned to show plausible behavior. The direct coupling from the TMS stimulation to the layer 5 cells is considerably lower than the indirect couplings from the layer 2/3 cells, reflecting the low amplitude of direct stimulation due to TMS deeper in the cortex. Time delays for the propagation to the layer 5 cells from the layer 2/3 cells are chosen to be consistent with Rusu et al. (2014); excitations far from the layer 5 soma requiring around 5 ms to travel to the soma. Others are commensurately quicker.

**Table A.1:** The standard parameter values used in the NFTsim model (Sanz-Leon et al., 2017) for the neural field modeling. The subscripts *e*, *i* and *v* denote layer 2/3 excitatory, layer 2/3 inhibitory and layer 5 excitatory corticospinal output neuron populations respectively. We have used the simplification that *γ*_*ab*_, *α*_*ab*_ and *β*_*ab*_ are dependent on only the presynaptic cell *b*; therefore only one subscript is used. The projections to the layer 5 cells are described with time delays *τ*_*vb*_ rather than axonal rate constants.

| Parameter | Description | Value | Unit |
| --- | --- | --- | --- |
| Excitatory cells, dendrite responses and propagation rates |  |  |  |
| $\alpha_e$ | dendritic rise rate | 280 | $s^{-1}$ |
| $\beta_e$ | dendritic fall rate | 70 | $s^{-1}$ |
| $\gamma_e$ | axonal rate | 110 | $s^{-1}$ |
| $\theta_e$ | Mean threshold for excitatory firing | $13 \times 10^{-3}$ | V |
| $\sigma_e$ | description of width of sigmoid distribution | $3.8 \times 10^{-3}$ | V |
| $Q_e^{\max}$ | Maximum excitatory firing rate | 340 | $s^{-1}$ |
| Inhibitory cells, dendrite responses and propagation rates |  |  |  |
| $\alpha_i^A$ | dendritic rise rate to GABA<sub>A</sub> input | 400 | $s^{-1}$ |
| $\beta_i^A$ | dendritic fall rate to GABA<sub>A</sub> input | 100 | $s^{-1}$ |
| $\alpha_i^B$ | dendritic rise rate to GABA<sub>B</sub> input | 40 | $s^{-1}$ |
| $\beta_i^B$ | dendritic fall rate to GABA<sub>B</sub> input | 10 | $s^{-1}$ |
| $\gamma_i$ | axonal rate | 1000 | $s^{-1}$ |
| $\theta_i$ | Mean threshold for inhibitory firing | $13 \times 10^{-3}$ | V |
| $\sigma_i$ | description of width of sigmoid distribution | $3.8 \times 10^{-3}$ | V |
| $Q_i^{\max}$ | Maximum inhibitory firing rate | 340 | $s^{-1}$ |
| Layer 5 cells |  |  |  |
| $\theta_v$ | Mean threshold for excitatory firing | $8 \times 10^{-3}$ | V |
| $\sigma_v$ | description of width of sigmoid distribution | $2.5 \times 10^{-3}$ | V |
| $Q_v^{\max}$ | Maximum excitatory firing rate | 900 | $s^{-1}$ |
| Synaptic couplings |  |  |  |
| $\nu_{ee}$ | e-to-e synaptic coupling | $1.92 \times 10^{-4}$ | V s |
| $\nu_{ei}^A$ | GABA<sub>A</sub>-modulated i-to-e synaptic coupling | $-0.72 \times 10^{-4}$ | V s |
| $\nu_{ei}^B$ | GABA<sub>B</sub>-modulated i-to-e synaptic coupling | $-0.72 \times 10^{-4}$ | V s |
| $\nu_{ie}$ | e-to-i synaptic coupling | $1.92 \times 10^{-4}$ | V s |
| $\nu_{ii}^A$ | GABA<sub>A</sub>-modulated i-to-i synaptic coupling | $-0.72 \times 10^{-4}$ | V s |
| $\nu_{ii}^B$ | GABA<sub>B</sub>-modulated i-to-i synaptic coupling | $-0.72 \times 10^{-4}$ | V s |
| $\nu_{ve}^{\text{fast}}$ | fast e-to-v synaptic coupling | $2.4 \times 10^{-4}$ | V s |
| $\nu_{ve}^{\text{slow}}$ | slow e-to-v synaptic coupling | $2.4 \times 10^{-4}$ | V s |
| $\nu_{vi}^A$ | GABA<sub>A</sub>-modulated i-to-v synaptic coupling | $-3.0 \times 10^{-4}$ | V s |
| $\nu_{vi}^B$ | GABA<sub>B</sub>-modulated i-to-v synaptic coupling | $-3.0 \times 10^{-4}$ | V s |
| Propagation delays |  |  |  |
| $\tau_{ve}^{\text{fast}}$ | fast e-to-v axonal delay<sup>36</sup> | $1 \times 10^{-3}$ | s |
| $\tau_{ve}^{\text{slow}}$ | slow e-to-v axonal delay | $5 \times 10^{-3}$ | s |
| $\tau_{ie}^A$ | GABA<sub>A</sub>-modulated i-to-v axonal delay | $3 \times 10^{-3}$ | s |
| $\tau_{ie}^B$ | GABA<sub>A</sub>-modulated i-to-v axonal delay | $3 \times 10^{-3}$ | s |

## Appendix B. Calcium dependent metaplasticity

The equations and parameters for calcium dependent metaplasticity are presented briefly here; for full details see (Fung and Robinson, 2014).

In the calcium-dependent plasticity (CaDP) scheme, the driver of plasticity is the postsynaptic intracellular calcium concentration [Ca^2+^]_*e*_ (where *e* represents the excitatory layer 2/3 population in the current work), modulated through NMDA receptors (Shouval et al., 2002). The ultimate excitatory-to-excitatory synaptic weight 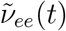 is modeled through (Fung and Robinson, 2014)

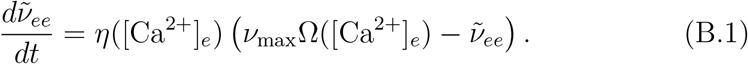

The 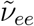 gives the value that the weight will ultimately come to when stimulation is stopped, thus it determines whether a protocol gives LTD or LTP. The parameter *ν*_*max*_ gives the maximum possible weight; Ω is dimensionless parameter that depends on [Ca^2+^]_*e*_ (0.5 for concentrations < 0.15 *μ*M, 0 for concentrations between 0.15–0.5 *μ*M and 1.0 at higher concentrations); *η* is a rate parameter that increases with increasing [Ca^2+^]_*e*_

The actual synaptic weight *ν*_*ee*_ responds slower and is modeled through

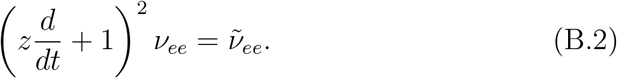

where *z* is a characteristic response timescale.

The postsynaptic calcium concentration [Ca^2+^]_*e*_ itself depends on the glutamate binding and postsynaptic activity through

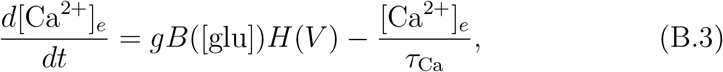

where *g* is the NMDA receptor-modulated calcium permeability, *B* is a sigmoidal function of the glutamate concentration [glu], *H* is voltage-dependent modulation of the dynamics (increasing with voltage *V* except at very high depolarizations), and *τ*_*Ca*_ a time-constant for calcium dynamics.

In the BCM approach to metaplasticity, the activity level that demarcates LTD from LTP is dependent on past activity. This is incorporated into CaDP by having conductance *g* dependent upon the history of the weight *ν*_*ee*_:

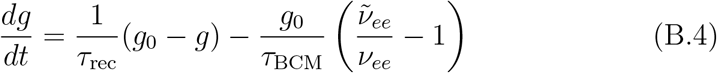

where *g*_0_ is the calcium conductance at equilibrium, *ν*_*ee*_ is the actual synaptic weight, and 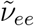 is the ultimate synaptic weight. The metaplasticity timescale is *τ*_*BCM*_ and the longer *τ*_*rec*_ is the recovery time for calcium conductance.

**Table B.2:** Parameters for the calcium dependent metaplasticity model, taken from Fung and Robinson (2013, 2014).

| Parameter | Description | Value | Unit |
| --- | --- | --- | --- |
| $\nu_{\max}$ | Maximum synaptic coupling | $10.0 \times 10^{-4}$ | V s |
| $\lambda_{\text{glu}}$ | Glutamate released per presynaptic excitatory spike | $50 \times 10^{-6}$ | M s |
| $g_0$ | NMDAR-modulated $\text{Ca}^{2+}$ conductance at equilibrium | $2 \times 10^{-3}$ | $\text{M s}^{-1} \text{ V}^{-1}$ |
| $\tau_{\text{Ca}}$ | Time-constant for calcium dynamics | $50 \times 10^{-3}$ | s |
| $\tau_{\text{glu}}$ | Time-constant for glutamate dynamics | $30 \times 10^{-3}$ | s |
| $\tau_{\text{rec}}$ | Timescale for recovery of calcium conductance | 1000 | s |
| $\tau_{\text{BCM}}$ | Timescale for metaplasticity | 7 | s |
| $z$ | Characteristic timescale for synaptic strength induction | 100 | s |

The glutamate concentration [glu] depends on presynaptic activity:

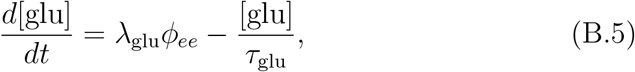

where *λ*_*glu*_ is the glutamate concentration released per presynaptic excitatory spike, *ϕ*_*ee*_ is the incoming synaptic from the excitatory population, Eq. (A.4), and *τ*_*glu*_ is the timescale for glutamate decay.

Equations (B.1) – (B.5), with functions *η*, Ω, *B* and *H* (Fung and Robinson, 2013), represent CaDP with metaplasticity. The resulting *ν*_*ee*_ is fed back into Eq. (A.2). Parameters and functions for these equations are as defined in Fung and Robinson (2013, 2014); parameters are listed in Table B.2.

## Appendix C. Response curves for different parameters

Figure C.9 shows the response curves for a range of different parameter sets. In each plot, the thick black line denotes the response for the standard parameter set, and the thin grey lines show the response when one parameter, indicated by the curve, has been changed by +15% (marked by the ‘+’ sign) or −15% (marked by the ‘−’ sign). The axonal delays, denoted by *τ*, are an exception. In these cases *τ* + refers to setting 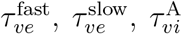 and 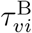 all to 3 ms, whereas *τ* − refers to setting 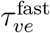 to 1 ms, 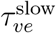 to 6 ms, and leaving 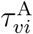 and 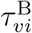 as 3 ms.

**Figure C.9:**
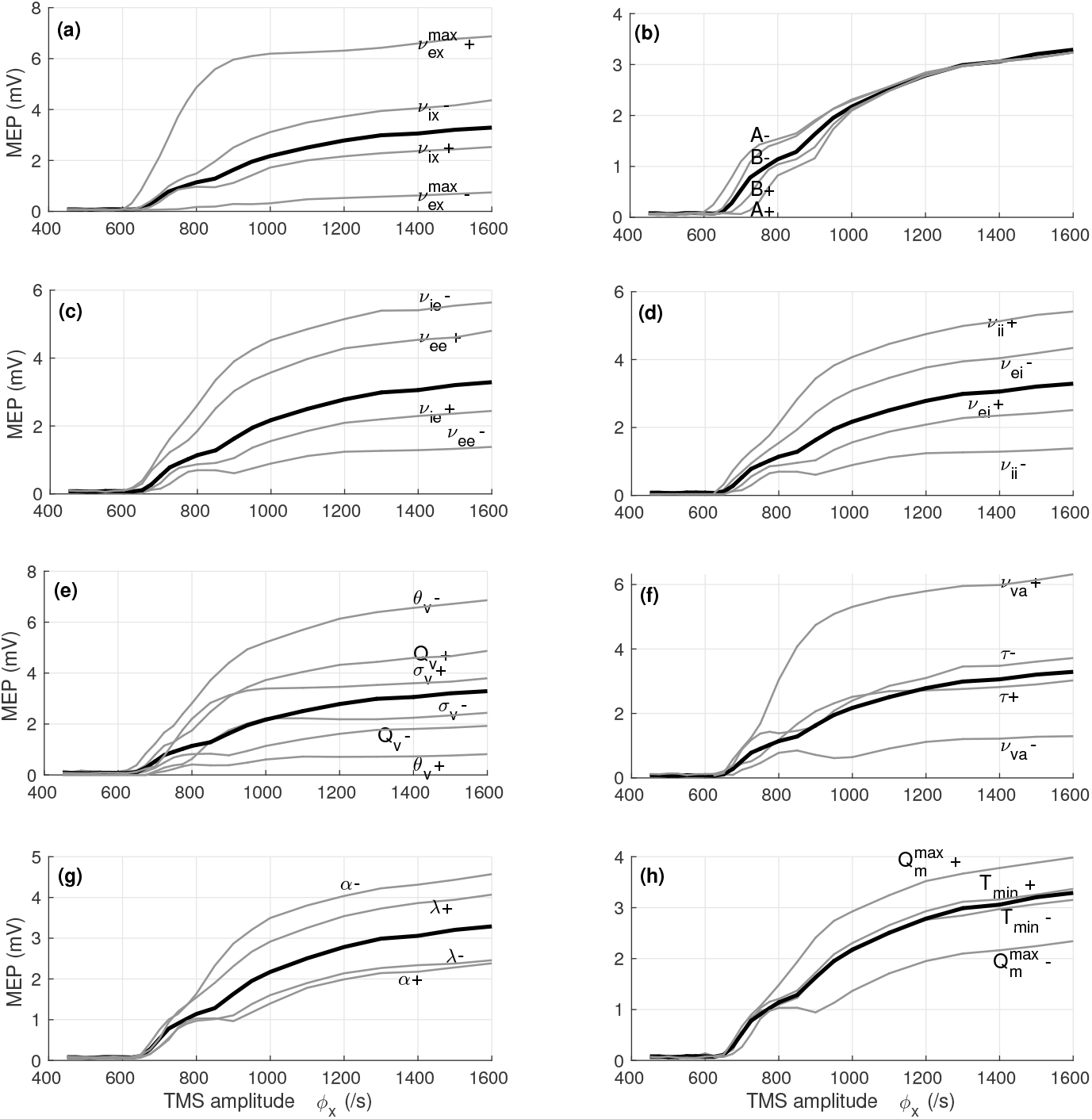
The motor evoked potential (MEP) as a function of external rate (TMS amplitude) *ϕ*_*x*_, for a variety of different parameter sets. For all plots, the thick black line shows the response for the standard parameter set of Appendix A. The thin gray lines show the effect of +15% changes (+) or 15% changes (−) in one (or sometimes more) of the parameters, as indicated. (a) and (b) The responses for changes in parameters associated with the TMS stimulation: maximum TMS-to-excitatory coupling 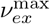; TMS-to-inhibitory coupling *ν*_*ix*_; TMS-to-excitatory coupling threshold *A*; width of TMS-to-excitatory coupling curve *B*. (c) Parameters describing coupling from excitatory cortical populations: excitatory-to-excitatory coupling *ν*_*ee*_; excitatory-to-inhibitory coupling *ν*_*ie*_. (d) Parameters describing coupling from inhibitory cortical populations: inhibitory-to-excitatory coupling *ν*_*ei*_; inhibitory-to-inhibitory coupling *ν*_*ii*_. (e) and (f) Parameters describing the corticospinal output neurons: layer 5 activation threshold *θ*_*v*_; maximum layer 5 firing rate 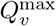; layer 5 activation curve width *σ*_*v*_; layer 2/3 to layer 5 couplings *ν*_*va*_; time delays for layer 2/3 to layer 5 *τ*. (g) and (h) Parameters describing the output neurons. exponent of firing threshold equation *α*; width of motor u4n0it action potential *λ*; maximum motoneuron firing rate 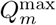; minimum threshold of motoneurons *T*_*min*_.

Table C.3 details the percentage changes in RMT, MEP at 200% RMT, and maximum gradient of the response curve, for the various parameter sets.

**Table C.3:** The effects of variation in model parameters on the response curve. For each change in parameters from baseline, percentage changes in resting motor threshold (RMT), motor-evoked potential (MEP) at 200% RMT (MEP_200_), and maximum gradient of the response curve are shown. For the axonal delay parameters *τ*, the listed cases are (i) setting 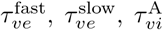 and 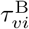 all to 3 ms (labeled ‘Equalized’) and (ii) setting 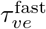 to 1 ms, 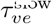 to 6 ms, and leaving 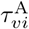 and 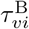 at 3 ms (labeled ‘Stretched’).

| Parameter | Description | Change | Changes in: |  |  |
| --- | --- | --- | --- | --- | --- |
|  |  |  | Threshold | MEP<sub>200</sub> | gradient |
| TMS stimulation |  |  |  |  |  |
| $\nu_{ex}^{\max}$ | Maximum TMS-to-$e$ coupling | +15% | -6% | 113% | 283% |
|  |  | -15% | 18% | -77% | -87% |
| $A$ | TMS-to-$e$ coupling threshold | +15% | 11% | 4% | -29% |
|  |  | -15% | -8% | -6% | 60% |
| $B$ | Width of TMS-to-$e$ coupling curve | +15% | 4% | 1% | 34% |
|  |  | -15% | -2% | -1% | 38% |
| $\nu_{vx}/\nu_{ex}$ | TMS-to-layer 5 : TMS-to-$e$ ratio | +15% | 0% | 0% | -2% |
|  |  | -15% | 0% | 0% | 3% |
| $\nu_{ix}$ | TMS-to-$i$ coupling | +15% | 2% | -23% | -25% |
|  |  | -15% | -2% | 30% | 45% |
| Layer 2/3 cells |  |  |  |  |  |
| $\nu_{ee}$ | $e$-to-$e$ coupling | +15% | -7% | 44% | 73% |
|  |  | -15% | 7% | -39% | -51% |
| $\nu_{ie}$ | $e$-to-$i$ coupling | +15% | 3% | -23% | -19% |
|  |  | -15% | -19% | 57% | -68% |
| $\nu_{ei}$ | $i$ to $e$ coupling | +15% | 2% | -25% | -38% |
|  |  | -15% | -2% | 31% | 22% |
| $\nu_{ii}$ | $i$ to $i$ coupling | +15% | -23% | 37% | -106% |
|  |  | -15% | 4% | -57% | -5% |
| Layer 5 cells |  |  |  |  |  |
| $\nu_{va}$ | layer 2/3 to layer 5 couplings | +15% | -1% | 99% | 105% |
|  |  | -15% | 0% | -60% | -43% |
| $\tau$ | $e$-to-layer 5 time delays | Equalized | 0% | -7% | -36% |
|  |  | Stretched | -1% | 19% | 151% |
| $Q_v^{\max}$ | Maximum layer 5 rate | +15% | -9% | 43% | 63% |
|  |  | -15% | 3% | -40% | -22% |
| $\theta_v$ | Layer 5 activation threshold | +15% | 10% | -75% | -93% |
|  |  | -15% | -27% | 62% | -103% |
| $\sigma_v$ | Layer 5 activation curve width | +15% | -4% | 9% | 32% |
|  |  | -15% | 8% | -24% | -32% |
| Motoneurons and muscle fibers |  |  |  |  |  |
| $T_{\min}$ | Minimum threshold | +15% | 0% | 5% | 33% |
|  |  | -15% | -10% | -10% | 15% |
| $\alpha$ | Exponent in Eq. (4) | +15% | 1% | -28% | 22% |
|  |  | -15% | -2% | 40% | 4% |
| $q$ | Minimum motoneuron firing rate | +15% | -2% | 1% | -9% |
|  |  | -15% | 1% | 0% | 50% |
| $Q_m^{\max}$ | Maximum motoneuron firing rate | +15% | -1% | 23% | 7% |
|  |  | -15% | 3% | -29% | 21% |
| $\lambda$ | Width of MUAP | +15% | -1% | 24% | 77% |
|  |  | -15% | 2% | -23% | -5% |

## Notes

### Competing Interest Statement

The authors have declared no competing interest.

### Summary of Updates

Resubmission to Clinical Neurophysiology following reviewer comments.

## References

Aberra, A.S., Wang, B., Grill, W.M., Peterchev, A.V.. Simulation of transcranial magnetic stimulation in head model with morphologically-realistic cortical neurons. Brain Stimulation 2020;13(1):175–189.

Breakspear, M.. Dynamic models of large-scale brain activity. Nature Neuroscience 2017;20:340–352.

Bungert, A., Antunes, A., Espenhahn, S., Thielscher, A.. Where does TMS stimulate the motor cortex? combining electrophysiological measurements and realistic field estimates to reveal the affected cortex position. Cerebral Cortex 2016;1–12.

Cooke, S.F., Bliss, T.V.P.. Plasticity in the human central nervous system. Brain 2006;129(7):1659–1673.

Day, B.L., Dressler, D., Maertens De Noordhout, A., Marsden, C.D., Nakashima, K., Rothwell, J.C., Thompson, P.D.. Electric and magnetic stimulation of human motor cortex: surface EMG and single motor unit responses. Journal of Physiology 1989;412:449–473.

Deco, G., Jirsa, V.K., Robinson, P.A., Breakspear, M., Friston, K.. The dynamic brain: from spiking neurons to neural masses and cortical fields. Public Library of Science Computational Biology 2008;4(8):e1000092.

Deng, Z.D., Lisanby, S.H., Peterchev, A.V.. Electric field depth-focality tradeoff in transcranial magnetic stimulation: Simulation comparison of 50 coil designs. Brain Stimulation 2013;6(1):1–13.

Devanne, H., Lavoie, B.A., Capaday, C.. Input-output properties and gain changes in the human corticospinal pathway. Experimental Brain Research 1997;114:329–338.

Di Lazzaro, V., Pilato, F., Dileone, M., Profice, P., Ranieri, F., Ricci, V., Bria, P., Tonali, P.A., Ziemann, U.. Segregating two inhibitory circuits in human motor cortex at the level of GABAA receptor subtypes: A TMS study. Clinical Neurophysiology 2007;118(10):2207–2214.

Di Lazzaro, V., Pilato, F., Dileone, M., Tonali, P.A., Ziemann, U.. Dissociated effects of diazepam and lorazepam on short-latency afferent inhibition. The Journal of Physiology 2005;569(1):315–323.

Di Lazzaro, V., Profice, P., Ranieri, F., Capone, F., Dileone, M., Olivero A. Pilato, F.. I-wave origin and modulation. Brain Stimulation 2012;5:512–525.

Di Lazzaro, V., Restuccia, D., Oliviero, A., Profice, P., Ferrara, L., Insola, A., Mazzone, P., Tonali, P., Rothwell, J.C.. Effects of voluntary contraction on descending volleys evoked by transcranial stimulation in conscious humans. The Journal of Physiology 1998;508(2):625–633.

Di Lazzaro, V., Rothwell, J.C.. Corticospinal activity evoked and modulated by non-invasive stimulation of the intact human motor cortex. The Journal of Physiology 2014;592(19):4115–4128.

Di Lazzaro, V., Ziemann, U., Lemon, R.N.. State of the art: Physiology of transcranial motor cortex stimulation. Brain Stimulation 2008;1(4):345–362.

Esser, S.K., Hill, S.L., Tononi, G.. Modeling the effects of transcranial magnetic stimulation on cortical circuits. Journal of Neurophysiology 2005;94(1):622–639.

Evarts, E.V.. Relation of pyramidal tract activity to force exerted during voluntary movement. Journal of Neurophysiology 1968;31:14–27.

Fung, P.K., Haber, A.L., Robinson, P.A.. Neural field theory of plasticity in the cerebral cortex. Journal of Theoretical Biology 2013;318:44–57.

Fung, P.K., Robinson, P.A.. Neural field theory of calcium dependent plasticity with applications to transcranial magnetic stimulation. Journal of Theoretical Biology 2013;324:72–83.

Fung, P.K., Robinson, P.A.. Neural field theory of synaptic metaplasticity with applications to theta burst stimulation. Journal of Theoretical Biology 2014;340:164–176.

Goetz, S.M., Alavi, S.M.M., Deng, Z., Peterchev, A.V.. Statistical model of motor-evoked potentials. IEEE Transactions on Neural Systems and Rehabilitation Engineering 2019;27(8):1539–1545.

Goldsworthy, M.R., Vallence, A.M., Hodyl, N.A., Semmler, J.G., Pitcher, J.B., Ridding, M.C.. Probing changes in corticospinal excitability following theta burst stimulation of the human primary motor cortex. Clinical Neurophysiology 2016;127:740–747.

Hallett, M.. Transcranial magnetic stimulation and the human brain. Nature 2000;406:147–150.

Hallett, M.. Transcranial magnetic stimulation: A primer. Neuron 2007;55:187–199.

Hamada, M., Murase, N., Hasan, A., Balaratnam, M., Rothwell, J.C.. The role of interneuron networks in driving human motor cortical plasticity. Cerebral Cortex 2013;23:1593–1605.

Hordacre, B., Goldsworthy, M.R., Vallence, A.M., Hamada, M., Rothwell, J.C., Ridding, M.C.. Variability in neural excitability and plasticity induction in the human cortex: A brain stimulation study. Brain Stimulation 2017;10:388–595.

Huang, Y.Z., Edwards, M.J., Rounis, E., Bhatia, K.P., Rothwell, J.. Theta burst stimulation of the human motor cortex. Neuron 2005;45:201–206.

Huang, Y.Z., Rothwell, J.C., Chen, R.S., Lu, C.S., Chuang, W.L.. The theoretical model of theta burst form of repetitive transcranial magnetic stimulation. Clinical Neurophysiology 2011;122(5):1011–1018.

Ilic, T.V., Meintzschel, F., Cleff, U., Ruge, D., Kessler, K.R., Ziemann, U.. Short-interval paired-pulse inhibition and facilitation of human motor cortex: the dimension of stimulus intensity. Journal of Physiology London 2002;545:153–167.

Inghilleri, M., Berardelli, A., Marchetti, P., Manfredi, M.. Effects of diazepam, baclofen and thiopental on the silent period evoked by transcranial magnetic stimulation in humans. Experimental Brain Research 1996;109(3):467–472.

Kiers, L., Cros, D., Chiappa, K.H., Fang, J.. Variability of motor potentials evoked by transcranial magnetic stimulation. Electroencephalography and Clinical Neurophysiology/Evoked Potentials Section 1993;89(6):415–423.

Kujirai, T., Caramia, M.D., Rothwell, J.C., Day, B.L., Thompson, P.D., Ferbert, A., Wroe, S., Asselman, P., Marsden, C.D.. Corticocortical inhibition in human motor cortex. Journal of Physiology 1993;471:501–519.

Laakso, I., Murakami, T., Hirata, A., Ugawa, Y.. Where and what TMS activates: Experiments and modeling. Brain Stimulation 2018;11:166–174.

Lefaucheur, J.P., André-Obadia, N., Antal, A., Ayache, S.S., Baeken, C., Benninger, D.H., Cantello, R.M., Cincotta, M., de Carvalho, M., De Ridder, D., et al. Evidence-based guidelines on the therapeutic use of repetitive transcranial magnetic stimulation (rtms). Clinical Neurophysiology 2014;125(11):2150–2206.

Li, B., Virtanen, J.P., Oeltermann, A., Schwarz, C., Giese, M.A., Ziemann, U., Benali, A.. Lifting the veil on the dynamics of neuronal activities evoked by transcranial magnetic stimulation. eLife 2017;6:e30552.

Li, X., Rymer, W.Z., Zhou, P.. A simulation based analysis of motor unit number index (MUNIX) technique using motoneuron pool and surface electromyogram models. IEEE Transactions in Neural Systems Rehabilitation Engineering 2012;20:297–304.

Liepert, J., Schwenkreis, P., Tegenthoff, M., Malin, J.P.. The glutamate antagonist riluzole suppresses intracortical facilitation. Journal of Neural Transmission 1997;104(11):1207–1214.

Matheson, N.A., Shemmell, J.B.H., De Ridder, D., Reynolds, J.N.J.. Understanding the effects of repetitive transcranial magnetic stimulation on neuronal circuits. Frontiers in Neural Circuits 2016;10:67.

McDonnell, M.N., Orekhov, Y., Ziemann, U.. The role of GABA(B) receptors in intracortical inhibition in the human motor cortex. Experimental Brain Research 2006;173(1):86–93.

Moezzi, B., Schaworonkow, N., Plogmacher, L., Goldsworthy, M.R., Hordacre, B., McDonnell, M.D., Iannella, N., Ridding, M.C., Triesch, J.. Simulation of electromyographic recordings following transcranial magnetic stimulation. Journal of Neurophysiology 2017;120:2532–2541.

Mohammadi, B., Krampfl, K., Petri, S., Bogdanova, D., Kossev, A., Bufler, J., Dengler, R.. Selective and nonselective benzodiazepine agonists have different effects on motor cortex excitability. Muscle & Nerve 2006;33(6):778–784.

Mori, F., Ribolsi, M., Kusayanagi, H., Siracusano, A., Mantovani, V., Marasco, E., Bernardi, G., Centonze, D.. Genetic variants of the NMDA receptor influence cortical excitability and plasticity in humans. Journal of Neurophysiology 2011;106(4):1637–1643.

Müller-Dahlhaus, J.F.M., Liu, Y., Ziemann, U.. Inhibitory circuits and the nature of their interactions in the human motor cortex a pharmacological TMS study. The Journal of Physiology 2008;586(2):495–514.

Murphy, S.C., Palmer, L.M., Nyffeler, T., Müri, R., Larkum, M.E.. Transcranial magnetic stimulation (tms) inhibits cortical dendrites. eLife 2016;5:e13598.

Olmo, G., Laterza, F., Lo Presti, L.. Matched wavelet approach in stretching analysis of electrically evoked surface EMG signal. Signal Processing 2000;80:671–684.

Opitz, A., Legon, W., Rowlands, A., Bickel, W.K., Paulus, W., Tyler, W.J.. Physiological observations validate finite element models for estimating subject-specific electric field distributions induced by transcranial magnetic stimulation of the human motor cortex. Neuroimage 2013;81:253–264.

Opitz, A., Windhoff, M., Heidemann, R.M., Turner, R., Thielscher, A.. How the brain tissue shapes the electric field induced by transcranial magnetic stimulation. NeuroImage 2011;58(3):849–859.

Parkin, B.L., Ekhtiari, H., Walsh, V.F.. Non-invasive human brain stimulation in cognitive neuroscience: a primer. Neuron 2015;87(5):932–945.

Pascual-Leone, A., Walsh, V., Rothwell, J.. Transcranial magnetic stimulation in cognitive neuroscience–virtual lesion, chronometry, and functional connectivity. Current opinion in neurobiology 2000;10(2):232–237.

Pashut, T., Magidov, D., Ben-Porat, H., Wolfus, S., Friedman, A., Perel, E., Lavidor, M., Bar-Gad, I., Yeshurun, Y., Korngreen, A.. Patch-clamp recordings of rat neurons from acute brain slices of the somatosensory cortex during magnetic stimulation. Frontiers in Cellular Neuroscience 2014;8:145.

Petreanu, L., Mao, T., Sternson, S.M., Svoboda, K.. The subcellular organization of neocortical excitatory connections. Nature 2009;457:1142–1145.

Pinotsis, D., Robinson, P., beim Graben, P., Friston, K.. Neural masses and fields: modeling the dynamics of brain activity. Frontiers in computational neuroscience 2014;8:149.

Reis, J., John, D., Heimeroth, A., Mueller, H.H., Oertel, W.H., Arndt, T., Rosenow, F.. Modulation of human motor cortex excitability by single doeses of amantadine. Neuropsychopharmacology 2006;31:2758–2766.

Roberts, J.A., Robinson, P.A.. Corticothalamic dynamics: Structure of parameter space, spectra, instabilities, and reduced model. Physical Review E 2012;85:011910.

Robinson, P., Rennie, C., Rowe, D., O’Connor, S.. Estimation of multiscale neurophysiologic parameters by electroencephalographic means. Human Brain Mapping 2004;23(1):53–72.

Robinson, P.A., Rennie, C.J., Wright, J.J.. Propagation and stability of waves of electrical activity in the cerebral cortex. Phys Rev E 1997;56:826–840.

Rogasch, N.C., Dartnall, T.J., Cirillo, J., Nordstrom, M.A., Semmler, J.G.. Corticomotor plasticity and learning of a ballistic thumb training task are diminished in older adults. Journal of Applied Physiology 2009;107(6):1874–1883.

Rogasch, N.C., Fitzgerald, P.B.. Assessing cortical network properties using TMS-EEG. Human Brain Mapping 2013;34(7):1652–1669.

Romero, M.C., Davare, M., Armendariz, M., Janssen, P.. Neural effects of transcranial magnetic stimulation at the single-cell level. Nature Communications 2019;10:2642.

Rusu, C.V., Murakami, M., Ziemann, U., Triesch, J.. A model of TMS-induced I-waves in motor cortex. Brain Stimulation 2014;7:401–414.

Sanz-Leon, P., Robinson, P.A., Knock, S.A., Drysdale, P.D., Abeysuriya, R.G., Fung, P.K., Rennie, C., Zhao, X.. NFTsim: Theory and simulation of multiscale neural field dynamics. PLoS Computational Biology 2017;14:e1006387.

Schwenkreis, P., Liepert, J., Witscher, K., Fischer, W., Weiller, C., Malin, J.P., Tegenthoff, M.. Riluzole suppresses motor cortex facilitation in correlation to its plasma level. Experimental Brain Research 2000;135(3):293–299.

Schwenkreis, P., Witscher, K., Janssen, F., Addo, A., Dertwinkel, R., Zenz, M., Malin, J.P., Tegenthoff, M.. Influence of the n-methyl-d-aspartate antagonist memantine on human motor cortex excitability. Neuroscience Letters 1999;270(3):137–140.

Seo, H., Jun, S.C.. Multi-scale computational models for electrical brain stimulation. Frontiers in Human Neuroscience 2017;11:515.

Shouval, H.Z., Bear, M.F., Cooper, L.N.. A unified model of NMDA receptor-dependent bidirectional synaptic plasticity. Proceedings of the National Acadamy of Sciences 2002;99:10831–10836.

Silva, S., Basser, P.J., Miranda, P.C.. Elucidating the mechanisms and loci of neuronal excitation by transcranial magnetic stimulation using a finite element model of a cortical sulcus. Clinical Neurophysiology 2008;119:2405–2413.

Tang, A.D., Lowe, A.D., Garrett, A.R., Woodward, R., Bennett, W., Canty, A.J., Garry, M.I., Hinder, M.R., Summers, J.J., Gersner, R., Rotenburg, A., Thickbroom, G., Walton, J., Rodger, J.. Construction and evaluation of rodent-specific rTMS coils. Frontiers in Neural Circuits 2016;10:47.

Thielscher, A., Opitz, A., Windhoff, M.. Impact of the gyral geometry on the electric field induced by transcranial magnetic stimulation. NeuroImage 2011;54(1):234–243.

Traub, R.D., Buhl, E.H., Gloveli, T., Whittington, M.A.. Fast rhythmic bursting can be induced in layer 2/3 cortical neurons by enhancing persistent na+ conductance or by blocking bk channels. Journal of Neurophysiology 2003;89(2):909–921.

Triesch, J., Zrenner, C., Ziemann, U.. Chapter 5 - modeling tms-induced i-waves in human motor cortex. In: Bestmann, S., editor. Computational Neurostimulation. Elsevier; volume 222 of Progress in Brain Research; 2015. p. 105–124.

Valls-Solé, J., Pascual-Leone, A., Wassermann, E.M., Hallett, M.. Human motor evoked responses to paired transcranial magnetic stimuli. Electroencephalography and Clinical Neurophysiology/Evoked Potentials Section 1992;85(6):355–364.

Wilson, M.T., Fulcher, B.D., Fung, P.K., Robinson, P.A., Fornito, A., Rogasch, N.C.. Biophysical modeling of neural plasticity induced by transcranial magnetic stimulation. Clinical Neurophysiology 2018;129:1230–1241.

Wilson, M.T., Fung, P.K., Robinson, P.A., Shemmell, J., Reynolds, J.N.J.. Calcium dependent plasticity applied to repetitive transcranial magnetic stimulation with a neural field model. Journal of Computational Neuroscience 2016;49:107–125.

Wilson, M.T., Goodwin, D.P., Brownjohn, P.W., Shemmell, J., Reynolds, J.N.J.. Numerical modelling of plasticity induced by transcranial magnetic stimulation. Journal of Computational Neuroscience 2014;36:499–514.

Wilson, M.T., Robinson, P.A., O’Neill, B., Steyn-Ross, D.A.. Complementarity of spike- and rate-based dynamics of neural systems. PLoS Computational Biology 2012;8:e1002560.

Wright, J.J., Liley, D.T.J.. Dynamics of the brain at global and microscopic scales: Neural networks and the EEG. Behavioral and Brain Sciences 1996;19:285295.

Zhou, P., Rymer, W.Z.. MUAP number estimates in surface EMG: Template-matching methods and their performance boundaries. Annals of Biomedical Engineering 2004;32:1007–1015.

Ziemann, U., Chen, R., Cohen, L.G., Hallett, M.. Dextromethorphan decreases the excitability of the human motor cortex. Neurology 1998;51:1320–1324.

Ziemann, U., Paulus, W., Nitsche, M.A., Pascual-Leone, A., Byblow, W.D., Berardelli, A., Siebner, H.R., Classen, J., Cohen, L.G., Rothwell, J.C.. Consensus: motor cortex plasticity protocols. Brain stimulation 2008;1(3):164–182.

Ziemann, U., Reis, J., Schwenkreis, P., Rosanova, M., Strafella, A., Badawy, R., Müller-Dahlhaus, F.. TMS and drugs revisited 2014. Clinical Neurophysiology 2015;126:1847–1868.

Ziemann, U., Rothwell, J.C., Ridding, M.C.. Interaction between intra-cortical inhibition and facilitation in human motor cortex. The Journal of physiology 1996;496(3):873–881.

Zrenner, C., Desideri, D., Belardinelli, P., Ziemann, U.. Real-time eeg-defined excitability states determine efficacy of tms-induced plasticity in human motor cortex. Brain Stimulation 2018;11(2):374–389.

